# Evaluating splicing factor and kinase network crosstalk through global phosphoproteomics

**DOI:** 10.64898/2026.04.16.718710

**Authors:** Sam Crowl, Sameek Singh, Tian Zhang, Kristen M. Naegle

## Abstract

Both splicing and kinase signaling are biochemical processes that fundamentally determine and shape cell physiology. Although there has been some indication that there is an interaction between the two – splicing can alter the availability of exons encoding kinase targets and kinases can phosphorylate splicing factors – it has yet to be established the extent to which altering splicing factor expression impacts kinase signaling networks. In this work, we implemented a data-driven analysis using ENCODE RNA-sequencing data and prior work mapping post-translational modifications onto splice events to identify candidate splice factor perturbations that show extensive alterations to phosphorylation-encoding protein products. We then replicated the ENCODE knockdown experiments and performed global phosphoproteomics for two candidates, U2AF1 and SRSF3, complementing the transcription-level data. Both knockdowns showed extensive changes in phosphorylation and kinase activities, both basally and upon receptor tyrosine kinase stimulation. U2AF1 knockdown drove decreased JNK-associated cell death signaling but elevated chromosome regulation through CSNK2A1, PLK, and EIF2AK4 activity. SRSF3 knockdown, on the other hand, led to decreased cell cycle signaling through CDK and HIPK2 but increased cytoskeletal signaling through various PAKs. In addition, we found a striking enrichment of phosphorylated splicing regulators in both knockdowns that were linked to their splicing activity, such as HNRNPC, suggesting potential feedback and crosstalk between splice factors through signaling pathway activation. Importantly, comparison of differential phosphorylation measurements from this study to mRNA expression and splicing measurements from ENCODE revealed significant knockdown-dependent protein regulation, not captured by transcriptomic measurements alone, underscoring the value of phosphoproteomic profiling after splice factor perturbations. Combined, the transcriptomics and phosphoproteomics reveal deep interconnection between the two processes that are relevant to understanding cell signaling in health and disease.

## Introduction

Alternative splicing is regulated by a complex network of RNA-binding proteins (RBP), called splice factors, which bind to pre-mRNA in a sequence- and context-dependent manner to direct exon inclusion and skipping [1]. Through the combinatorial action of multiple factors, cells can precisely control which isoforms are produced in response to developmental, tissue-specific, or environmental cues [1, 2]. Signaling pathway activity frequently modulates the activity, localization, and RNA-binding properties of splicing regulators, but there are also an increasing number of examples of alternative splicing remodeling signaling networks by altering the isoform composition and structure of key pathway components [1–4]. Alternative exons often encode for intrinsically disordered regions, short linear motifs, and phosphorylation sites, raising the possibility that splice factor expression can reshape kinase networks across tissues and disease contexts [5–7]. In our prior work, for example, we highlighted coordinated regulation of serum glucocorticoid kinase (SGK) signaling through ESRP1-dependent splicing of SGK substrates in prostate cancer [7]. Similarly, NOVA-dependent splicing in neurons was shown to preferentially impact kinases and phosphorylation-encoding regions [8]. Further, kinase isoform-specific signaling outcomes have been observed in diverse contexts. For example, the FynT isoform of the SRC family kinase FYN promotes inflammatory signaling and is upregulated in Alzheimer’s disease, while the brain-predominant FYN isoform (FynB) is upregulated in various hematological malignancies [9–11]. Splicing can even introduce unexpected signaling behaviors, such as a PKBFB4 isoform found in hepatocellular carcinoma that phosphorylates AKT [12]. Altogether, there is strong evidence to suggest that splice factor expression and activity can facilitate widespread changes to global phosphorylation and kinase networks across different tissue and disease contexts. Despite this, there is limited available data that directly connects RNA splicing perturbations to phosphorylation at a systems-level.

In this work, we wished to more directly evaluate the impact of splice factor knockdown on phosphorylation and establish a phosphoproteomic complement to pre-existing transcriptomic measurements. For this, we first harnessed comprehensive measurements of splice factor binding preferences and knockdown effects on mRNA from ENCODE [13]. With our prior approach to identify PTMs associated with splice events and their downstream effects (PTM-POSE [7]), we mapped the effects of knockdown-associated exon inclusion on phosphorylated protein regions and RTK signaling to identify candidate splice factors with the strong potential to induce changes to phosphorylation networks through mRNA splicing. Using this data-driven approach, we selected two candidate splice factor knockdowns (U2AF1 and SRSF3), repeated ENCODE-project knockdowns, and performed global phosphoproteomics. We found that knockdown of either factor led to widespread global phosphorylation changes – in fact, phosphorylation differences were a greater function of splice factor knockdown than broad RTK-stimulated effects, suggesting splice factor alterations create significant alteration to kinase signaling network context. Following global analysis of the phosphoproteome, we integrated mRNA data from ENCODE with the phosphoproteomic complement to understand the possible ways in which splice factors and kinases interact at the network level. While some phosphorylation differences after knockdown could be linked to transcript-level alterations in expression or splicing, many could not be explained by direct mRNA regulation, suggesting substantial indirect effects. We also found similar phosphorylation observed after both SRSF3 and U2AF1 knockdown, suggesting some degree of shared response to knockdown, including the phosphorylation of many other splice factors. Overall, this study provides one of the first profiles of the impact of splice factor knockdown on global phosphorylation and kinase activity. Given the prevalence of splice factor mutations and activity in diseases like cancer, establishing clear connections between splice factor regulation and signaling pathways could reveal new biomarkers or avenues for therapies. In moving towards this integrated understanding, this study lays out methods for RBP selection and subsequent interpretation of mRNA and phosphorylation data as it relates to the interaction of splicing and phospho-signaling.

## Results

### Systems-level characterization of RNA-binding protein effects on phosphorylation

ENCODE provides a rich resource characterizing the transcriptional response of cells to RNA binding protein (RBP) knockdown [13]. In order to evaluate the possible connections of splice factors to alterations in available phosphorylation sites, we used PTM-POSE [7], to map post-translational modifications to ENCODE RNA-sequencing data for knockdowns where the RBP is a splice factor based on UniProtKB definitions (61 total knockdowns) [13]. We quantified differential splicing upon knockdown using rMATS, filtered for significant splice events, and identified phosphorylation sites associated with each splice event using PTM-POSE [7, 14]. Knockdowns resulted in highly variable differences in the number of splice events (ranging from 121 to 4,053 significant splice events) and impacts on phosphorylation-encoding regions (ranging from 5.7% to 28.8% of splice events linked to phosphorylation-encoding regions, Figure S1A).

Focusing on the RBP knockdowns that resulted in a large proportion of phosphorylation-encoding splice events, we next evaluated splice event associations using NEASE pathway enrichment [15]. Out of this combined analysis, we selected two candidate splice factors for secondary experimental analysis to explore phosphorylation effects directly – U2AF1 and SRSF3. Both factors are associated with a higher percentage of splice events associated with phosphorylation-encoding regions (25.2% and 27.7%, respectively for U2AF1 and SRSF3; Figure S1B). Prior work has indicated that SRSF3’s preference for cytosine-rich motifs leads to inclusion of exons associated with serine and threonine residues, which can be phosphorylated [16]. Additionally, NEASE analysis suggested both U2AF1 and SRSF3 knockdown were associated with splice events enriched for proteins involved in RTK signaling (Figure S1C), supporting the likelihood that splicing effects on phosphorylation and signaling proteins indicates intentional functional roles of these splice factors. Notably, both U2AF1 and SRSF3 have been found to be altered in cancers, with U2AF1 being mutated in lung cancers and acute myeloid leukemia and SRSF3 being upregulated in various cancers, increasing their disease relevance [3, 17, 18]. Thus, we chose to focus on the impacts of the U2AF1 and SRSF3 knockdown on the global phosphoproteome and RTK signaling, based on this data-driven analysis of possible phosphorylation effects, broad regulatory connections to RTK signaling, and relevance to disease.

Next, we wished to measure the direct phosphoproteomic effect of U2AF1 and SRSF3 knockdown, complementing ENCODE transcriptomic data. Using ENCODE protocols, we performed knockdown in HepG2 cells, where we validated shRNA knockdown via western blot, finding greater than 50% decrease in signal for both knockdowns (Figure S2, Figure S3), relative to a GFP-targeting shRNA control. Since our goal was to ensure active signaling and phosphorylation in the network, we included a serum starved and “RTK-activated” treatment, where we stimulated with a mix of HepG2-relevant growth factors (EGF, HGF, and insulin). We performed RTK-stimulation briefly (5-minutes), which is ample time to evoke tyrosine and serine/threonine signaling cascades, but before substantial transcriptomic effects have occurred, helping to ensure our phosphoproteomic data is best matched to ENCODE-based transcriptomic profiling. We confirmed RTK-stimulation resulted in sufficient signaling activation by western-based monitoring of phosphorylated ERK and AKT, key signaling nodes in the transition from RTK-mediated phosphotyrosine networks to downstream transcriptional, cytoskeleton, and metabolic outcomes (Figure S2, Figure S3).

Having recreated successful knockdown of our target splice factors in HepG2 cells and established an approach to increase kinase network activation, we performed phosphoproteomic analysis in triplicate, across the three knockdown conditions (shGFP, shU2AF1, shSRSF3), with both pre- and post-RTK activation. We identified a total of 24,818 phosphorylation sites across 5,485 genes that were captured in at least two replicates, with ∼43% of sites found in all conditions, allowing for cross-condition comparison (Figure 1B). Principal component analysis (PCA) demonstrated that the phosphoproteomic data had strong replicate similarity, suggesting high reproducibility of this discovery-based phosphoproteomic approach (Figure 1C). We first evaluated phosphorylation on the perturbed splice factors, finding, in line with its expected expression loss, SRSF3 observed decreased phosphorylation at T24 after knockdown (no U2AF1 phosphorylation sites were observed). Next, in order to confirm RTK activation resulted in physiologically expected outcomes, we explored activation-specific phosphorylation sites in each condition (greater than 2-fold increases with statistical significance; Figure S5A). Some RTK-activated sites were shared across all knockdown conditions, including sites in several genes known to be involved in early RTK signaling like GAB1 and CRKL (Table S2). Next, we predicted kinase activity based on using KSTAR [19] with the phosphoproteomic data. This analysis found activation of many of the kinases associated with early RTK signaling, including MET, EGFR/ERBB2, AKT, and ERK (Figure S5B). Hence, global phosphoproteomics demonstrated highly reproducible measurements and significant coverage of RTK-activated kinase networks.

**Figure 1.**
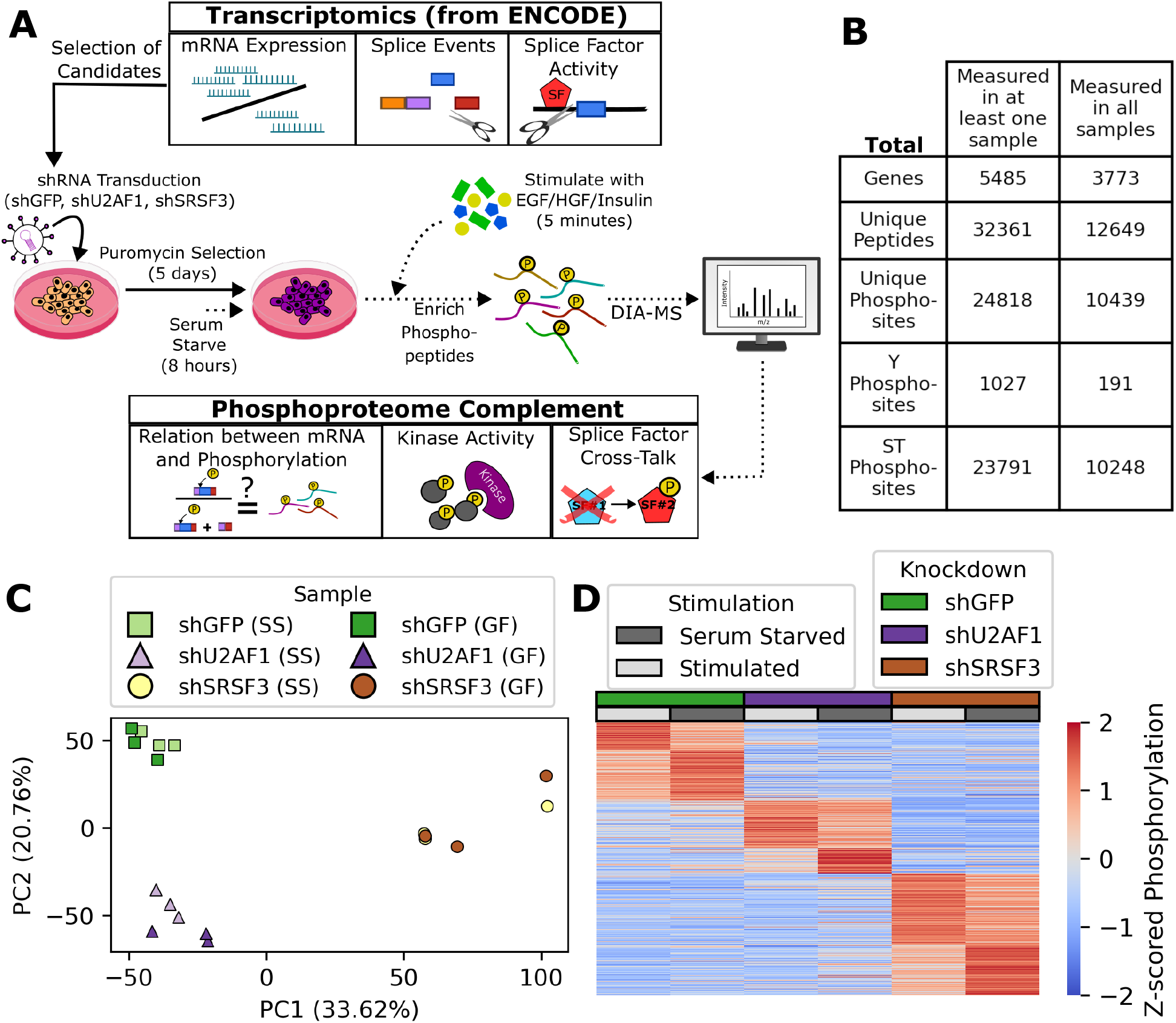
Using and complementing transcriptomic approaches of splice factor knockdown to explore effects on the phosphoproteome. **A)** We used ENCODE-based transcriptomic data to identify RNA binding protein candidates that have widespread effects on kinase signaling. We then replicated knockdown in HepG2 and performed the phosphoproteomic analysis of two selected candidates (U2AF1 and SRSF3) either as serum starved or short term stimulated with a cocktail of growth factors to elicit early kinase network signaling. A suite of analysis tools were applied to both the transcriptomic data from ENCODE and the complement of phosphoproteomic data. **B)** The number of unique genes, peptides and sites measured by global phosphoproteomics. Peptides are only considered if covered in at least two of the three replicates in a condition. **C)** Principal component analysis (PCA) of measured phosphopeptides in each replicate, for peptides captured in all eighteen replicates. **D)** Comparison of average phosphopeptide measurements for each experimental condition. Replicates from the same experimental condition were averaged, and then the mean intensities were standardized across conditions via Z-score normalization. Peptides were sorted by a two-step K-means clustering, where three clusters were first found, which were then each broken into two subclusters.

One facet of this data that emerged robustly is that RBP knockdown substantially altered the phosphoproteome of cells. This can be seen in the weak correlation between U2AF1 and SRSF3 RTK-regulated sites (*r* = 0.2; Figure S5C), with very few common genes or sites showing RTK-based regulation (only 67 shared sites, Figure S5D,E). Globally, we observed from PCA that the phosphoproteomic differences were largest between knockdown conditions, rather than between serum starvation and RTK-activation. Clustering-based analysis of the phosphoproteomic data reinforces this, identifying three distinct groupings that are knockdown-specific, with secondary groups that capture RTK stimulation (Figure 1D). Hence, this phosphoproteomic data suggests that RBP knockdown has the capacity to significantly reshape the phosphorylation landscape of the cell, both its basal and activated states.

Finally, since we selected U2AF1 and SRSF3 based on data-driven analysis of possible splicing effects on phosphorylation through transcriptomic-based analysis (i.e. alterations in inclusion and exclusion of phosphorylation-containing exons), we returned to analyze the coverage of splicing-associated phosphorylation sites from ENCODE in the complementary phosphoproteomic data. Within each condition, we measured between 442 and 556 phosphotyrosine sites and 15,554 and 16,935 phosphoserine sites by mass spectrometry (MS). Based on ENCODE data and rMATS splice events, a small fraction (< 5%) of the sites measured in the phosphoproteomic dataset were associated with splice events perturbed by U2AF1 or SRSF3 knockdown (Figure S4A). To validate prior findings that splicing changes were correlated with phosphorylation abundance (i.e. the larger proportion of transcripts with phosphorylation-encoding exon, the likelier phosphorylation on the protein was seen), we compared fold changes in phosphorylation after SRSF3 or U2AF1 knockdown relative to control to changes in isoform inclusion quantified from ENCODE data. In line with our prior work, we found a weak to moderate correlation (r = 0.23 - 0.39) between splicing inclusion changes and phosphorylation abundance (Figure S4C). However, it should be noted that this analysis was limited by the coverage of phosphorylation sites we expected to be impacted by U2AF1/SRSF3 splicing. Of the 6,834 phosphorylation sites associated with either U2AF1 or SRSF3 splice events from ENCODE data, we were only able to obtain measurements in our phosphoproteomic dataset for 944 of these sites (13.8%), which was particularly notable for tyrosine sites (3.6% and 5.3% of U2AF1- and SRSF3-associated sites were measured by MS, respectively; Figure S4B). Overall, the data-driven analysis identified both RBP knockdowns had substantial effects on the phosphorylation across kinase signaling networks, but that direct splicing of phosphorylation-containing exons explains only a small fraction of the widespread changes observed under U2AF1 and SRSF3 knockdown.

### Evaluating splice factor-specific and shared effects on kinase signaling

Global phosphoproteomics demonstrated profound phosphorylation network effects across the cell as a result of splice factor knockdown, including distinct differences in patterns between the two splice factors. Our next objective was to explore the potential biological consequence of phosphorylation network changes and their connection to kinase signaling and regulation. Given that we observed similar phosphorylation trends after knockdown in pre- and post-RTK stimulation conditions (Figure S6), we combined pre- and post-RTK stimulation replicates for further analysis to capture the holistic effect of splicing factor knockdown on the phosphorylation footprint. To identify kinases relevant to each knockdown, we used several tools for kinase activity prediction (KSTAR [19], KSEA [20], and Kinase Library [21]), as each approach has different coverage of kinases and different approaches for using the phosphoproteomic data to predict kinase activity impacts. We also identified enriched functions of differentially phosphorylated sites (based on PhosphoSitePlus site-specific annotations [22]) and proteins (based on gene sets from Reactome [23]) across RBP-specific and shared effects. Finally, we utilized the available transcriptomic data from ENCODE to contextualize differential phosphorylation and kinase activity changes as it relates to transcriptional and splicing effects of U2AF1 and SRSF3 knockdown.

#### Effects on the phosphoproteome following U2AF1 knockdown

U2AF1 is a core auxiliary factor of the U2 spliceosome important for early initiation of pre-mRNA spicing, interacting with many other splicing regulators including U2AF2, SF1, SF3B1, and even SRSF3 [24, 25]. U2AF1 knockdown, relative to GFP control, resulted in 3,580 differentially phosphorylated sites and was fairly balanced between both up- and down-regulated sites (51% which increase and 49% of sites decrease; Figure 2A). Based on annotations from PhosphoSitePlus, differentially phosphorylated sites tend to be associated with cell cycle regulation, DNA repair, and endocytosis (Figure S7A). Genes with increasing phosphorylation were enriched for cell cycle and DNA/RNA regulation pathways such as nonhomolgous end-joining (NHEJ) (Figure 2B). Interestingly, genes with decreasing phosphorylation were more associated with GTPase and cell death signaling. This included depleted phosphorylation of genes associated with JNK-associated signaling through NGAGE, including a 14-3-3 interaction site in ARHGEF2 (S960) and a KRAS interaction site in NGEF (S63). Holistically, this suggests U2AF1 knockdown drives increasing phosphorylation on mitotic genes and decreases in cell death signaling, painting a potentially pro-survival signaling phenotype in HepG2 cells. Notably, there have been conflicting reports on whether U2AF1 expression is pro- or anti-proliferative depending on the cell context, although U2AF1 is consistently connected to cell cycle and mitotic genes in these studies [18, 26, 27].

**Figure 2.**
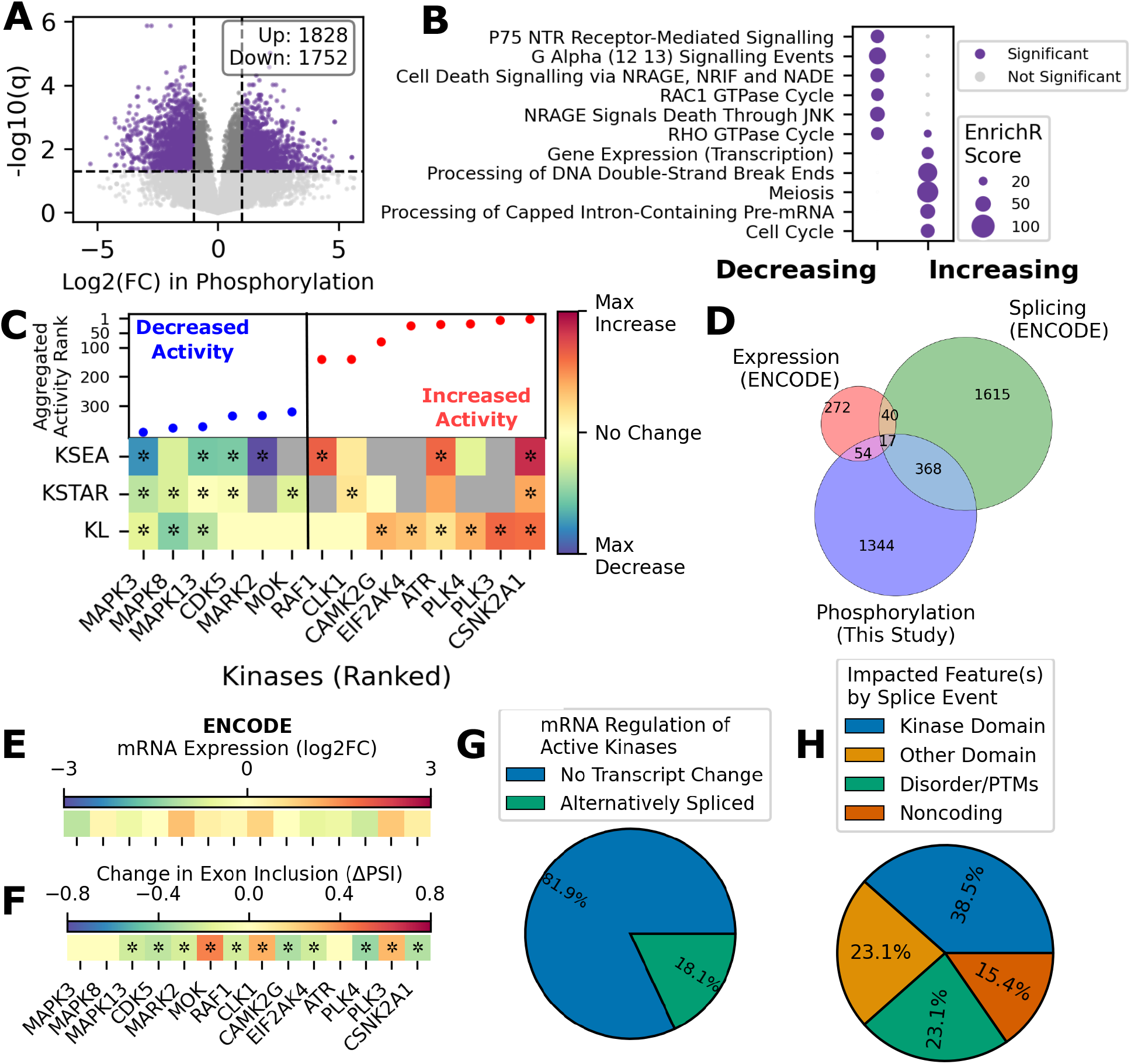
Profiling the perturbed phosphoproteome after U2AF1 knockdown. **A)** We identified sites with significantly increasing or decreasing phosphorylation after U2AF1 knockdown relative to the GFP control, considering samples under serum starvation and RTK-activation (*q* ≤ 0.05, |*LFC*| ≥ 1). Statistical significance was assessed with a two-tailed t-test and corrected using Benjamini-Hochberg FDR correction. **B)** Enriched gene sets among differentially phosphorylated genes, based on Reactome pathway gene sets [23] and calculated using EnrichR [28]. **C)** Kinase activity ranks (a subset are shown) from three substrate-based tools (KSEA [20], KSTAR [19], KinaseLibrary (KL) [21]). The aggregated activity rank and direction of significance is indicated in the top panel, with kinase activity changes predicted from each tool shown in the bottom panel, with significant activities denoted with * (KSEA: *p* ≤ 0.05, KSTAR: *FPR*_*up or down*_ ≤ 0.05, KL: *p*_*adj,up or down*_ ≤ 0.001). Full plot with all kinases in Figure S9 and Table S4. **D)** Number of genes that were differentially phosphorylated after U2AF1 knockdown in this study compared to the genes identified to be alternatively spliced or have differential expression based on ENCODE data [13]. **E)** Log2 fold changes in mRNA expression for the subset of kinases shown in Panel C, based on ENCODE data. *: *p*_*adj*_ ≤ 0.05. **F)** Change in exon inclusion for the subset of kinases shown in Panel C, based on rMATS analysis of ENCODE data. *: *FPR* ≤ 0.05. **G)** For kinases with differential activity (Panel C), we identified the fraction that exhibited changes in the kinase’s mRNA (expression, splicing). **H)** For kinases with predicted differential activity and associated with an alternative splice event, we identified the transcript or protein features impacted by the splicing of each kinase, including domains, PTMs, and noncoding regions.

We next identified kinases that are predicted to change in activity following knockdown (Figure 2C, Figure S9, Table S4). Across all three activity prediction approaches, we found 43 kinases with increasing activity and 28 kinases with decreasing activity. There was a significant increase in activity of CSNK2A1 after knockdown, associated with elevated phosphorylation of various transcription and DNA repair genes like XPC S94, NCOA2 S469, and DNMT3A S390. Similarly, we observed increased activity of stress associated pathways through PLK3/4 and EIF2AK4, where PLK3/4 in particular has been connected to DNA repair signaling. On the other hand, we observed decreased activity of the stress-activated kinase MAPK8 (JNK1), matching gene set enrichment results connecting decreasing phosphorylation to NGAGE-associated JNK signaling. There was also decreased activity of other MAP kinases, including MAPK3 (ERK1) and MAPK13, which match the slightly depleted phosphorylation of MAPK1/3 after knockdown measured by western blot (Figure S3). Finally, CDK5, a mediator of cytoskeletal dynamics and p53 apoptotic signaling, also observed decreased activity after knockdown. Altogether, kinase activity was broadly impacted by knockdown, including increasing and decreasing activities. Interestingly, U2AF1 knockdown appears to cause a distinct switch in the activated stress signaling pathways, moving away from JNK and CDK5 stress signaling associated with programmed cell death and towards DNA repair and mitosis pathways mediated by PLK and CSNK2A1.

Next, we wished to evaluate the connection between mRNA-level regulation and phosphorylation using RNA-sequencing measurements from ENCODE. In line with U2AF1’s predominant role as a splicing regulator, the majority of mRNA-regulation of transcripts was through splicing – 2,041 genes were alternatively spliced, whereas only 383 genes were differentially expressed, based on ENCODE data (Figure 2D, Figure S8A). There is a small degree of overlap in genes regulated at the mRNA-level with their protein products that are differentially phosphorylated – 385 of the spliced genes (18.9%) and 71 of the differentially expressed genes (18.5%) were differentially phosphorylated. In these instances, changes to mRNA expression or splicing were weakly to moderately related to phosphorylation changes (Figure S4C, Figure S8B). However, most phosphorylation differences (75.3%) were not directly connected to changes in mRNA regulation (Figure 2D). Given this, we next looked for mRNA expression or splice events that may explain the phosphorylation differences after U2AF1 knockdown (Figure 2E,F), focusing on mRNA-based regulation of kinases. Across the 71 kinases with predicted differential activity, 13 underwent a significant splice event (18.05%; Figure 2G) and none had significant mRNA expression changes (Figure 2G, Figure S8A). For those 13 kinases that were differentially spliced, we evaluated splicing impacts on protein features (Figure 2H). We found a variety of splice effects, including splicing of the kinase domain (CDK5, MAPK13, CLK1/2, and MOK), binding domains (like the Polo-box domains of PLK3/4), PTM-encoding regions (NEK1, RAF1, MARK2), and even some noncoding regions (CSNK2A1, CAMK2G). Hence, for at least some kinases, there appears to be a direct mechanistic link between alternative splicing and activity, which then accounts for some aspect of the change in phosphorylation states in the network. However, the vast majority of kinase activity changes, and the phosphorylation events connected to them, cannot be connected directly to mRNA regulation, suggesting that there is a complex interconnection between splicing and kinase networks.

#### Effects on the phosphoproteome following SRSF3 knockdown

SRSF3 is a member of the SR protein family that binds to exonic splicing enhancer (ESEs) elements to facilitate splice site selection by the spliceosome (including the U2 spliceosome). Although SRSF3 is predominantly known for its role in regulating splicing, it has also been connected to transcription regulation and mRNA stability [29]. SRSF3 knockdown, relative to GFP control, results in 6,721 differentially phosphorylated sites, fairly balanced between both up- and down-regulated sites (51.9% which increase and 48.1% of sites decrease) (Figure 3A). Interestingly, this split in up and down regulation is consistent with U2AF1. However, SRSF3-knockdown produces almost twice the number of differentially phosphorylated events than U2AF1. Similar to U2AF1 knockdown, many of these sites tend to be associated with cell cycle regulation and DNA repair, but we also found an enrichment of phosphorylation sites important for regulating the cytoskeleton and apoptosis (Figure S7B). When profiling the differentially phosphorylated genes, we found GTPase signaling was generally enriched among differential phosphorylation events (Figure 3B). Interestingly, genes associated with chromatin modifications and epigenetic regulation were less phosphorylated (such as transcription repressors like NCOR1/2), while genes associated with mRNA processing and splicing were elevated in phosphorylation (Figure 3B). Prior transcriptomic analyses [30] suggested that SRSF3 knockdown led to mRNA regulation of genes associated with cell growth, splicing, and cytoskeletal regulation, suggesting that similar biological pathways are regulated at both the transcriptional and post-translational level.

**Figure 3.**
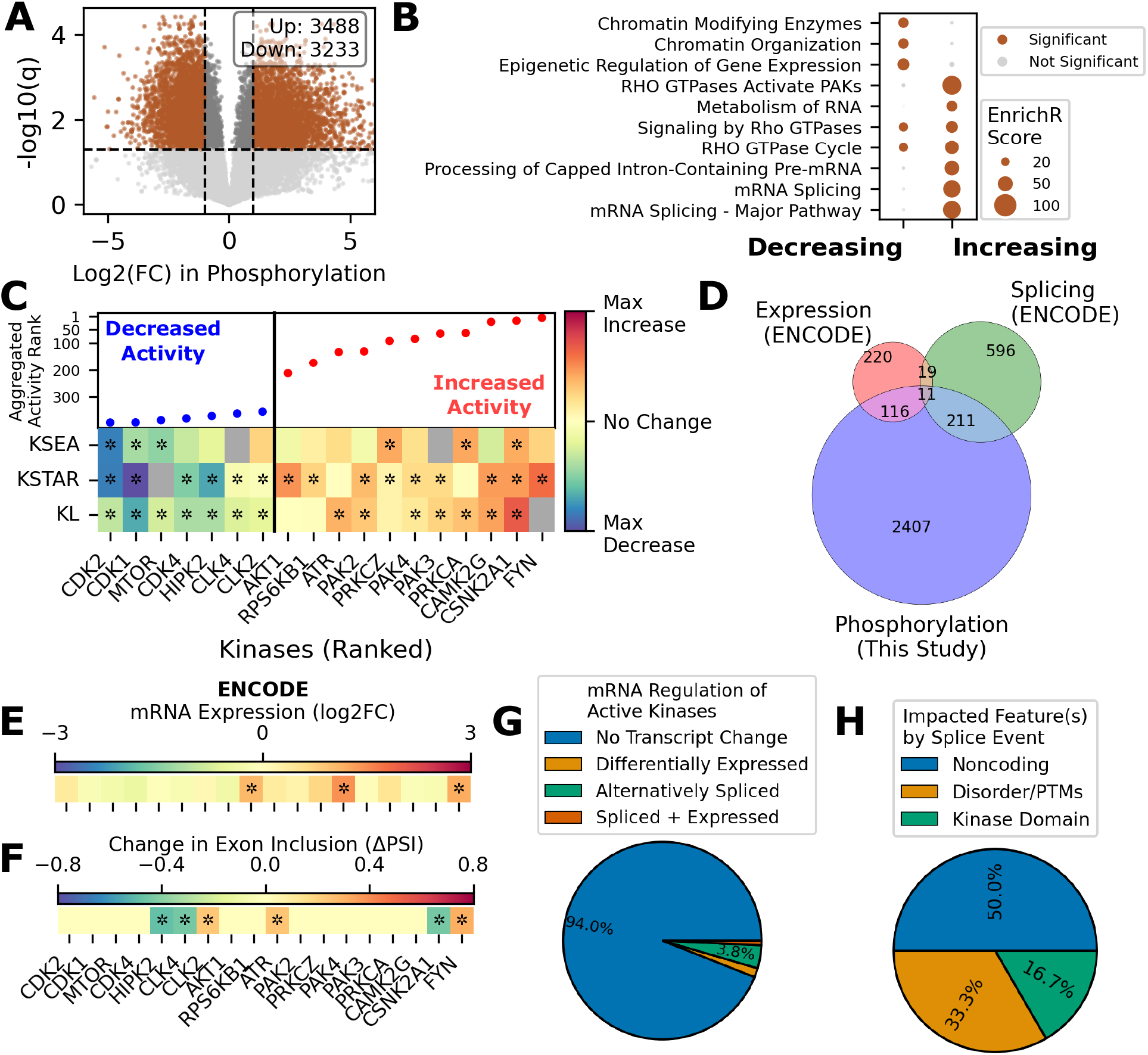
Profiling the perturbed phosphoproteome after SRSF3 knockdown. **A)** We identified sites with significantly increasing or decreasing phosphorylation after SRSF3 knockdown relative to the GFP control, considering samples under serum starvation and RTK-activation (*q* ≤ 0.05, |*LFC*| ≥ 1). Statistical significance was assessed with a two-tailed t-test and corrected using Benjamini-Hochberg FDR correction. **B)** Enriched gene sets among differentially phosphorylated genes, based on Reactome pathway gene sets [23] and calculated using EnrichR [28]. **C)** Kinase activity ranks and differences from three substrate-based tools (KSEA [20], KSTAR [19], KinaseLibrary (KL) [21]) for a subset of kinases measured. The aggregated activity rank and direction of significance is indicated in the top panel, with kinase activity changes predicted from each tool shown in the bottom panel, with significant activities denoted with * (KSEA: *p* ≤ 0.05, KSTAR: *FPR*_*up or down*_ ≤ 0.05, KL: *p*_*adj,up or down*_ ≤ 0.001). Full plot with all kinases in Figure S9 and Table S4. **D)** Number of genes that were differentially phosphorylated after SRSF3 knockdown in this study compared to the genes identified to be alternatively spliced or have differential expression based on ENCODE data [13]. **E)** Log2 fold changes in mRNA expression for the subset of kinases shown in Panel C, based on ENCODE data. *: *p*_*adj*_ ≤ 0.05. **F)** Change in exon inclusion for the subset of kinases shown in Panel C, based on rMATS analysis of ENCODE data. *: *FPR* ≤ 0.05. **G)** For kinases with differential activity (Panel C), we identified the fraction that exhibited changes in the kinase’s mRNA (expression, splicing). **H)** For kinases with predicted differential activity and associated with an alternative splice event, we identified the transcript or protein features impacted by the splicing of each kinase, including domains, PTMs, and noncoding regions

Next, we identified a total of 100 kinases with increasing activity and 33 kinases with decreasing activity based on at least one activity inference tool (Figure 3C, Figure S9B, Table S4). In line with prior reports of SRSF3-knockdown induced cell cycle arrest and cellular senescence [31], we found decreased activity of cell cycle kinases CDK1/2/4 as well as MTOR and HIPK2. Among kinases with increasing activity, there were several PI3K-AKT pathway associated kinases, including AKT1, RPS6KB1, PRKACA, and FYN, which may explain the elevated phosphorylation of AKT observed from western blots after RTK activation (Figure S2). In addition, matching the gene set enrichment results, we found increased activity of several PAK kinases (PAK2/3/4), which are best known for their role in cytoskeletal regulation and dynamics [32]. Altogether, kinase activity analysis highlights widespread inhibition of cell-cycle related signaling but upregulation of cell motility and cytoskeletal regulation pathways, explaining the decreased cell proliferation and altered morphology we qualitatively observed after SRSF3 knockdown.

When comparing differential phosphorylation patterns with mRNA regulation, we similarly found a small degree of overlapping regulation (222 and 127 differentially phosphorylated genes were alternatively spliced or differentially expressed, respectively). However, 2,042 differentially phosphorylated genes (87.7%) were not perturbed at the RNA level (Figure 3D). We were able to connect some of the differentially active kinases to changes in mRNA regulation (Figure 3E,F), including several kinases that increased in mRNA expression and had elevated kinase activity after knockdown (RPS6KB1, PAK4, and FYN). Just as in U2AF1 knockdown, CSNK2A1 undergoes a splice event in the noncoding region associated with elevated activity (Figure 3F, Figure S10A,B). We also found alternative splice events in HIPK2, CLK2, and CLK4 associated with decreased activity, while splice events in ATR and FYN were associated with increased activity. Of the six alternatively spliced kinases with differential activity (3.8%), three of the splice events impacted noncoding regions (CSNK2A1, ATR, CLK4), one impacted the kinase domain and disordered regions (FYN), and two impacted disordered and PTM-encoding regions (HIPK2, CLK2) (Figure 3G,H). Notably, the splice event in HIPK2 was previously validated after SRSF3 knockdown and is associated with an E3 ligase binding region, leading to decreased proteosomal degradation after knockdown [31]. However, although it was originally suggested to not impact activity level of HIPK2, our global analysis suggest decreased activity of HIPK2, indicating additional functions of the spliced HIPK2 region related to phosphorylation of transcription-related genes like NCOR1, SUPT20H, and NCOA6 (Figure S10C,D). However, the vast majority of differentially active kinases (95%) could not be connected to either mRNA expression or splicing changes, an even smaller percentage than observed for U2AF1 knockdown (Figure 3G).

#### Shared regulation between U2AF1 and SRSF3 phosphorylation effects

Given that we observed some degree of kinase activities between U2AF1 and SRSF3 knockdown (such as elevated activity of CSNK2A1), we were curious whether there were similarities between the phosphorylation profiles after U2AF1 and SRSF3 knockdown. Interestingly, we found a moderate correlation between fold changes in phosphorylation after U2AF1 and SRSF3 knockdown (*r* = 0.53; Figure 4A). Nearly half of the significantly regulated sites were shared between both knockdowns, with 732 sites increasing and 681 sites decreasing after knockdown (Figure 4A). There were also 288 sites with opposing regulation (increasing in one knockdown and decreasing in the other), many of which were associated with cell cycle regulation like RB1 T821 and CDK Y15. Based on this, we binned the knockdown-associated sites into four main categories – SRSF3-specific changes, U2AF1-specific changes, broad splicing perturbations (shared by both U2AF1 and SRSF3), and opposing regulation (opposite direction of regulation in both knockdowns). Based on PhosphoSitePlus annotations, phosphorylation sites associated with both knockdowns were broadly associated with cell cycle regulation, transcription, and cell motility (Figure 4B). SRSF3-specific sites were uniquely associated with apoptosis, cell adhesion, and autophagy, while U2AF1-specific sites were uniquely enriched for sites associated with chromatin organization and endocytosis. When assessing specific biological processes associated with differentially phosphorylated genes, we found similar patterns of splice-factor specific regulation of chromatin organization (U2AF1) and cell adhesion/cadherin binding (SRSF3) (Figure 4C, Table S3). Further, we found a striking association with splicing regulation and RNA processing among the genes with increasing phosphorylation after both U2AF1- and SRSF3-knockdown, which may indicate phosphorylation-mediated feedback and crosstalk between splice factors previously only observed via mRNA regulation.

**Figure 4.**
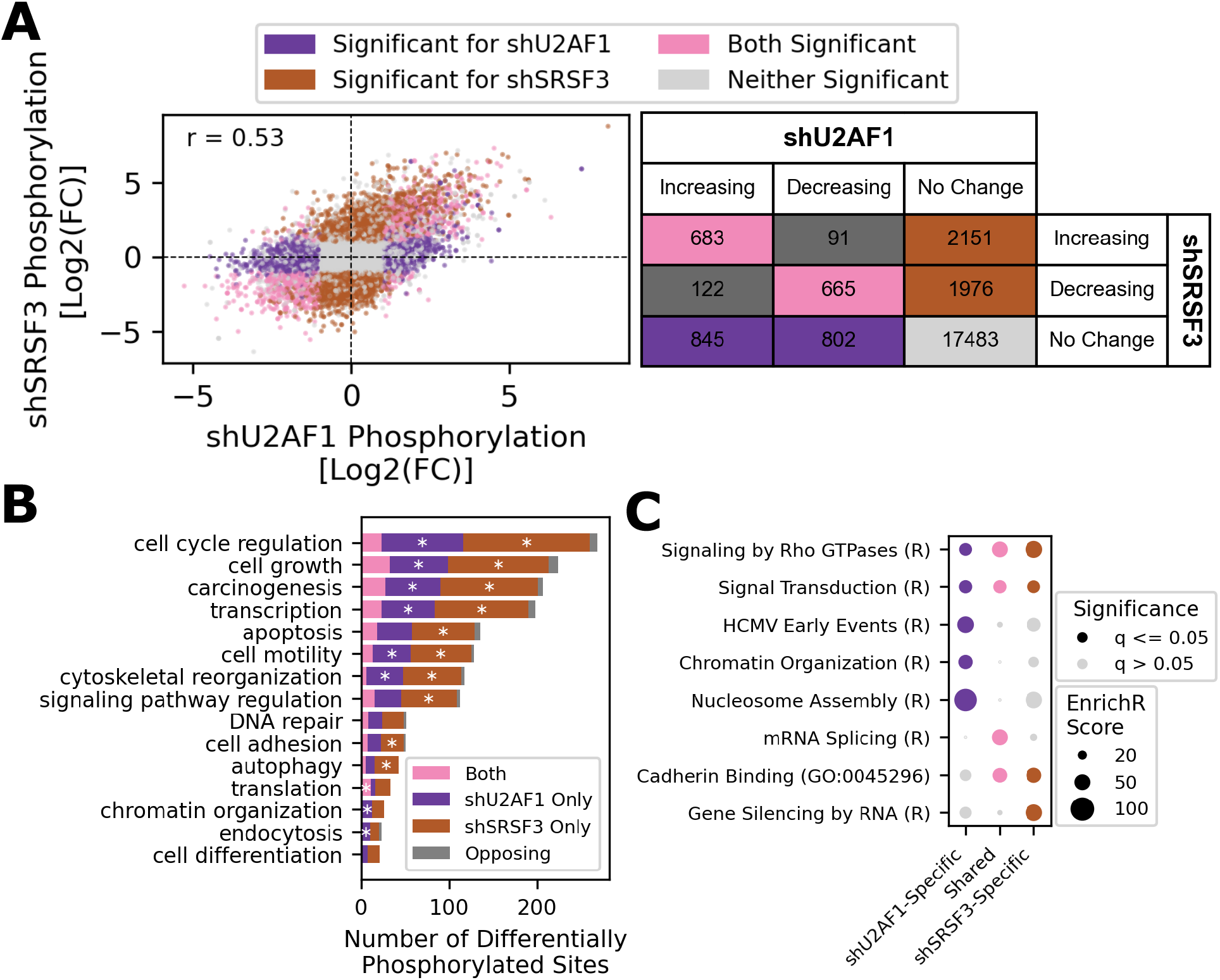
Comparing the shU2AF1 and shSRSF3 phosphoproteomes. We identified differentially phosphorylated sites after U2AF1 and SRSF3 shRNA knockdown relative to the shGFP control, considering perturbations under both serum starvation and stimulation conditions. **A)** Comparison of phosphorylation site fold changes between U2AF1 and SRSF3 knockdown, relative to GFP control. Points are colored by whether the fold change is statistically significant in shU2AF1 (purple), hSRSF3 (brown), or both conditions (pink), based on a two-tailed t-test and Benjamini-Hochberg FDR correction (*q* ≤ 0.05). The table on the right side of the plot shows the counts within each group (U2AF1-specific, SRSF3-specific, or both). **B)** Biological processes associated with differentially phosphorylated sites unique to U2AF1 or SRSF3 knockdown, shared by both, or with conflicting directions across knock-downs. Site-specific annotations were obtained from PhosphoSitePlus [22]. Enrichment was assessed using a Fisher’s exact test, using all sites measured in phosphoproteomic dataset as the background (*: *q* ≤ 0.05). **C)** Enrichment of Reactome(R)/Gene Ontology (GO) gene sets among differentially phosphorylated genes associated with U2AF1 knockdown only (purple), SRSF3 knockdown only (brown), or both knockdowns (pink).

### Exploring the mechanisms connecting kinase signaling and splice factor knockdowns

Following phosphoproteomic complementation of splicing factor knockdown, it is clear that there are substantial and far-reaching effects on signaling networks, reshaping both their basal state and the response capabilities. In our analysis of knockdown effects on phosphorylation, we found some of those network alterations involve direct regulation of kinase activity and that kinases are phosphorylating splice factors. We next wished to explore more deeply evidence for the direct regulation of kinases by splicing factors and the regulation of splicing factors by kinases, enabled by combining the transcriptomic and phosphoproteomic measurements.

#### Transcriptional regulation of kinases and its connection to kinase activity

Given the primary role of U2AF1 and SRSF3 in mRNA regulation and the observation that several alternatively spliced kinases had perturbed kinase activities, we were curious the extent to which post-transcriptional regulation of kinases is predictive of kinase activity changes. Of the 18 differentially expressed kinases after SRSF3 or U2AF1 knockdown based on ENCODE data, we found 25-50% were connected to a kinase with differential activity, including FYN and RPS6KB1, after SRSF3 knockdown (Figure S11C,D). Across the 73 splice events affecting kinases with measured activity across either knockdown, approximately 25% were associated with a significant activity change (Figure S11C,D). We characterized the types of spliced protein regions to determine if specific types of splice events may be more likely to influence a kinase’s activity (domains, PTMs, etc.). Although there were no significant associations with specific feature types and activity, splice events that impacted regions encoding PTMs produced the highest rate of altered activity, while splice events impacting domains were no more likely to influence activity than other splice events (Figure S11). Of the 19 splice events affecting an internal region of the kinase domain, 4 were correlated with loss of activity (CDK5, CLK2, MAPK13, and MOK), which may indicate a regulatory mechanism to influence kinase activity without altering expression for these kinases. For example, loss of a 35aa insert in the CDK5 kinase domain was associated with loss of activity in U2AF1 knockdowns, an isoform that has previously been identified as being expressed and localized in the nucleus of skeletal muscle and the testis (Figure S10E,F) [33]. In addition to coding regions, we also found two examples of differentially active kinases undergoing a splice event in noncoding regions, important for transcript stability and translation (CSNK2A1 in both knockdowns, CAMK2G in U2AF1 knockdown) (Figure S10A,B). Ultimately, these examples highlight how RBP perturbations may directly influence kinase activity, but also given significant alterations in kinase activity beyond splicing, highlights the deep interconnection in kinase networks, for which activity cannot be described directly by transcriptional control.

#### Transcriptional and kinase regulation of splice factor activity

Phosphorylation plays a prominent role in the regulation of splice factor localization, binding, and splicing activity [2]. Given the observed enrichment of splicing-related genes among differentially phosphorylated genes, we sought to better characterize the phosphorylation of splicing regulators and its potential consequence after both U2AF1 and SRSF3 knockdown (Figure 5A). We found 99 and 130 splicing-related RBPs that were differentially phosphorylated in response to U2AF1 and SRSF3 knockdown, respectively, comprising nearly half of the splicing-related RBPs assessed (264 total). Phosphorylation of these RBPs was strongly correlated between knockdowns, even more so than across the overall phosphoproteome (*r* = 0.72; Figure 5A), where 74.6% of the sites were shared as significantly different in both conditions (Figure 5A). Whereas generally phosphorylation was seen to be evenly split between increasing and decreasing, on RBPs there is a skew towards increasing phosphorylation following U2AF1 and SRSF3 knockdown. Since phosphorylation of RNA-binding domains (RBDs) has previously been shown to alter binding specificity to RNA [], we evaluated splicing factor phosphorylation location within the RNA binding domains, finding that majority of phosphorylation events fall outside of domains, with only ∼7% of regulated phosphorylation sites occurring within RBDs (Figure 5B).

**Figure 5.**
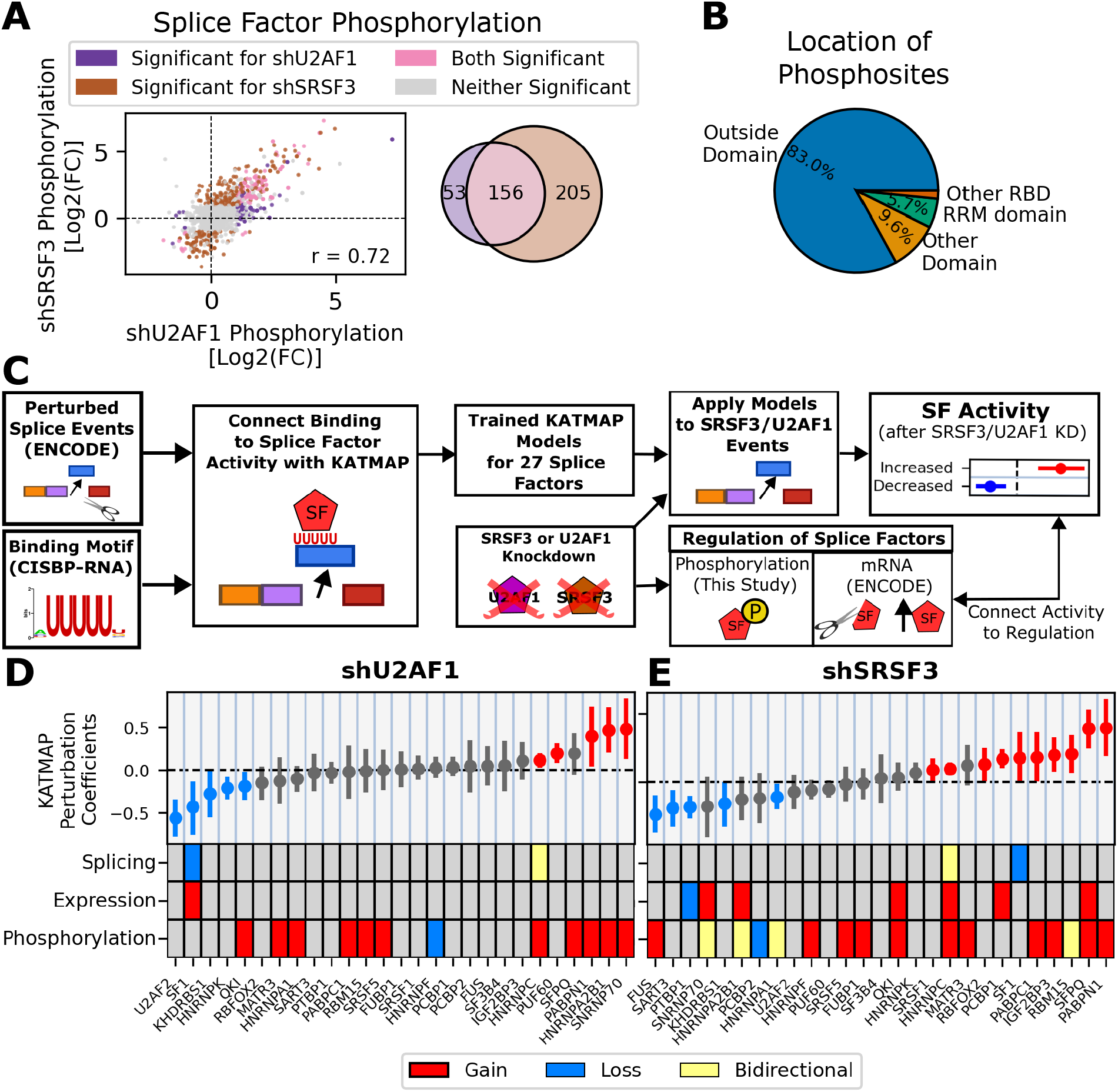
Phosphorylation of splice factors and their relationship to splicing activity. We characterized the phosphorylation and activity of splice factors after knockdown. **A)** Comparison of the phosphorylation differences of 246 splicing-related RBPs (based on UniProt/ENCODE annotations) after U2AF1 or SRSF3 knockdown. Left panel shows comparison between fold changes for individual phosphorylation sites, with the venn diagram illustrating the number of significantly differentially phosphorylated sites after U2AF1 and SRSF3 knockdown. Color indicates statistical significance for shU2AF1 (purple), shSRSF3 (brown), or both conditions (pink) based on a two-tailed t-test and Benjamini Hochberg FDR correction (*q* ≤ 0.05). **B)** Fraction of differentially phosphorylated sites after either U2AF1 or SRSF3 knockdown residing in RNA-binding domains (RBD) such as the RRM domain, other domains, or are outside domains. **C)** Pipeline to generate 27 KATMAP models of splice factor activity and apply them to U2AF1 or SRSF3-associated splice events. We compared changes to splice factor activity to mRNA regulation from ENCODE (expression, splicing) or phosphorylation from this study. **D**,**E)** KATMAP model predictions for U2AF1 knockdown (D) or SRSF3 knockdown (E) are shown in the top panel, where red indicates significant increases in splicing activity and blue indicates significant decreases in splicing activity. Direct splicing factor regulation occurring after knockdown (mRNA expression/splicing from ENCODE, phosphorylation from this study) is highlighted in the bottom panels, including the direction of regulation. Expression and phosphorylation significance defined as: |*Log*2*FC*| ≥ 0.5 and *q* ≤ 0.05. Splicing significance is defined as: |Δ*PSI*| ≥ 0.2 and *FDR* ≤ 0.05.

To better understand the overall network of splice factors, and how kinases may play a role in regulating that network, we harnessed KATMAP, a tool for predicting the splice factor activity from RNA sequencing measurements after a perturbation [34]. Using ENCODE knockdown data for 27 different splice factors and corresponding RNA binding motifs from CISBP-RNA [13, 35], we generated trained KATMAP models that connect splice factor binding to splice event regulation for each splice factor. We then used the trained models to predict splice factor activity in the U2AF1 and SRSF3 knockdown conditions. Unfortunately, we were not able to generate a sufficient model for either U2AF1 or SRSF3 splicing activity (see methods for details). Based on KATMAP model predictions, U2AF1-associated splicing was connected to 5 splice factors with increasing activity and 5 with decreasing activity. SRSF3-associated splicing was connected to 10 splice factors with increasing activity and 5 with decreasing activity (Figure 5C). As positive validation of this approach, U2AF2, a key interaction partner of U2AF1, was predicted to have decreased splicing activity only in U2AF1 knockdown, indicating the expected disruption of normal U2 spliceosome function [**?**]. In general, despite the similarities in phosphorylation and splicing profiles, the splice factors predicted to be influencing alternative splice events after each knockdown were mostly unique, with only three commonalities – increased activity of HNRNPC and PABPN1 and decreased activity of KHDRBS1.

Next, to contextualize splice factor activities, we evaluated possible regulatory mechanisms of splice factors with altered activity, including mRNA expression, splicing changes, and phosphorylation (Figure 5D). Phosphorylation changes on these splice factors was more substantial than mRNA-based alterations. For the splice factors that show coordinated activity changes in both knockdowns, KHDRBS1 was not associated with any clear mRNA processing or phosphorylation changes, but both HNRNPC (S260) and PABN1 (S19) exhibited elevated phosphorylation (Figure S12). Notably, HNRNPC S260 was previously noted to influence mRNA binding and alter HNRNPC splicing activity. In many cases, differences in phosphorylation patterns between the knockdowns were associated with distinct splicing activity. This was most prominent for cases where U2AF1 and SRSF3 led to opposing phosphorylation of RBPs, such as HNRNPA2B1, HNRNPA1, SNRNP70, and RBM15. One prominent example of this is SNRNP70, a key component of the U1 spliceosome which was predicted to have significantly increased activity after U2AF1 knockdown and decreased (although not significant) activity after SRSF3 knockdown (Figure 5D,E). Of note, Y126 (in the RRM domain) and S226 (previously identified to induce splicing through interaction with snRNPs [36]) were depleted in SRSF3 knockdown, while Y126 phosphorylation was elevated after U2AF1 knockdown, suggesting that phosphorylation of these sites modulate activity of SNRNP70 and are perturbed differently in each knockdown (Figure S12). Similar patterns can be observed for other splice factors, like IGF2BP3 being phosphorylated at S438 and residing in one of its KH domains after SRSF3 knockdown, correlating with elevated activity (Figure S12). Notably, many of the splice factors with affected activity did not exhibit altered mRNA expression or splicing based on ENCODE data, meaning it is missed by the traditional transcriptomic measurements from ENCODE. Additionally, not all phosphorylation events could be connected to a change in splice factor activity, (e.g. SRSF5 and MATR3 were similarly phosphorylated in both knockdowns but had no predicted change in splicing activity), suggesting that interpreting phosphoproteomic effects also requires understanding transcriptional-level regulation.

## Discussion

There have been strong connections between splicing and kinase signaling throughout recent research, with both of these biochemical processes being important to cell determination and response to external stimuli. Splice factors serve as central mediators of this interconnectedness, as they are key regulators of splicing decisions and are themselves phosphorylated. This study is the first of its kind to systematically query the extent of splice factors’ importance to kinase signaling. Even though we selected specifically for splice factors that would likely have an effect on the overall kinase signaling network, we were surprised at the sheer extent of change in the global phosphorylation network capabilities – where the same cell line (HepG2) under the same stimulation are so completely different from both the control and from each other. Activity analysis highlights the reshaping of network configurations by splice factors and kinases occurring after knockdown, as well as a strong relationship between the two biochemical processes and their regulatory networks. Importantly, transcriptomic data alone provides an incomplete picture of the functional consequence of splice factor expression. By profiling the global phosphorylation footprint in tandem with transcriptomic data, we can generate more informed hypotheses about the functional consequences of splice factor knockdown. For example, we identified 51 kinases with potentially relevant alternative splice events after U2AF1 knockdown based on ENCODE datasets, most of which were not associated with any known functional impact. Without phosphoproteomic data and subsequent kinase activity profiling of the knockdown data, it would be challenging to isolate which splice events have the largest impact on kinase networks. With phosphoproteomic data, we were able to narrow this list to 13 high-priority kinases that warrant further investigation moving forward.

Perhaps most strikingly, widespread phosphorylation of splice factors themselves revealed signaling consequences that are entirely invisible to transcriptomic approaches. For example, CSNK2A1 is known to phosphorylate HNRNPC, which exhibited increasing splicing activity after both U2AF1 and SRSF3 knockdown. In this way, there was a connection between elevated CSNK2A1 and HNRNPC activity after both U2AF1 and SRSF3 knockdown, which may be explained a splice event in the noncoding region of CSNK2A1, elevated activity of CSNK2A1, and subsequent increased phosphorylation of HNRPNC at S260. Similarly, CDK2 and CLKs showed decreased activity after SRSF3 knockdown and have been shown to phosphorylate SNRNP70 at S226, which was depleted after SRSF3 knockdown but not after U2AF1 knockdown. As with CSNK2A1, both CLK2/4 undergo splice events after SRSF3 knockdown based on ENCODE data, highlighting a potential mechanism by which phosphorylation of SNRNP70 is altered. While further validation is needed, these findings highlight the utility of a multiomic approach to studying splice factor function and indicate that signaling consequences of splice factor expression are broader than transcriptomic measurements imply.

It should be noted that this study was not designed to capture the dynamics of splicing regulation, which misses information about the primary splice events that lead to initial phosphorylation changes, subsequent splicing activity, and further kinase alterations resulting from network feedback. A dynamic study could eventually help disentangle the waves of regulation in the two networks arising from both the direct and indirect consequences of splice factor expression, an important consideration given the self-regulating and combinatorial nature of both kinase and splice factor networks. In addition, although this discovery-based phosphoproteomic approach helped lay important groundwork for understanding the role of splice factors in altering phosphorylation networks, by using a global approach, we missed key phosphotyrosine events that would further establish RTK-specific network alterations. Future targeted approaches might be helpful in understanding more specifically regulation sites across all conditions. Finally, given the investment needed to go from knockdown to phosphoproteomic measurement, we did not test if the data-driven approach was capable of identifying splice factor perturbations that had large versus small effects on kinase signaling networks, instead focusing on characterizing those that likely had effects. It would be an excellent follow on study to identify whether there are splice factors who do not significantly reshape kinase signaling networks, which would give the field a better understanding of how to think about alterations in specific splice factors in health and disease.

While we were not focused on a specific disease context in this study, aberrant regulation of both alternative splicing and kinase signaling are common contributors to disease genesis and progression, with kinases and splice factors commonly exhibiting genomic alterations across various cancers (amplifications, mutations, etc.). [3, 37]. The two splice factors examined here, SRSF3 and U2AF1, are themselves recurrently altered or overexpressed in various cancers, making the signaling consequences we observed particularly relevant to disease contexts []. Dysregulated signaling in cancer commonly drives alternative splicing alterations through phosphorylation of downstream splice factors, and conversely, aberrant splicing can reshape kinase networks [4, 38]. This is a bidirectional relationship our data directly illustrates. Further, alternative splicing has been suggested to be a driver of drug resistance, such as SRSF7-dependent splicing of ATM driving resistance to chemotherapy or a truncated form of HER2 that promotes resistance to trastuzumab, a HER2 targeting antibody [39, 40]. For this reason, while kinase inhibitors are one of the most common targets of FDA-approved drugs for oncology, there is increasing interest in developing therapies targeting the spliceosome and associated splice factors [3]. This includes the potential for combination strategies with kinase inhibitors, such as dual targeting of SF3B1 and BTK in chronic lymphocytic leukemia (CLL) cells [41]. Given their connection to cell signaling regulation, splice factors could also potentially serve as biomarkers for kinase inhibitor efficacy. However, there remains a paucity of data exploring the functional impact of splice factor activity on signaling outcomes, with only one recent study profiling the impact of spliceosome inhibition on the proteome [42]. By profiling the impact of U2AF1 and SRSF3 knockdown on the phosphoproteome, we highlighted the complex interplay between splice factors and kinases that goes overlooked when only transcript regulation is assessed. Future analyses could build on this framework to explore splice factor activity in different disease contexts, including in cancer models with splice factor mutations, specific kinase alterations, or under kinase inhibition.

## Methods

### Quantifying splice events associated with RBP knockdown experiments from ENCODE

To assess the impact of RBP-mediated splicing on kinases and cell signaling, we utilized RBP knockdown experiments from the ENCODE database, focusing on experiments in HepG2 cells [13]. To prioritize alterations from alternative splicing, we restricted analysis to RBPs associated with the ‘RNA-binding’ and ‘mRNA splicing’ keywords in UniProt [43]. In total, we analyzed 61 different RBP-knockdown experiments. For each experiment, we first downloaded bam alignment files (GencodeV29) from ENCODE using the REST API. In addition, differential expression was downloaded for the same experiments and annotation set. We quantified splice events occurring after knockdown using rMATS-turbo (v4.1.1) and the downloaded bam files [14]. We specified the library type as ‘fr-firststranded’, read length as 100, and allowed for variable read length. Finally, PTMs associated with significant splice events were identified using PTM-POSE v0.3.1 [7]. Unless otherwise specified, we focused on splice events (and associated PTMs) with *FDR*≤ 0.05, Δ*PSI* ≥ 20%, and with a minimum of 20 junction or exon reads for both the knockdown and control samples.

For each significant splice event after SRSF3 and U2AF1 knockdown, we identified the exons and potential transcripts associated with the event based on genomic information from Ensembl [44]. We then annotated each splice event with the relevant protein features (PTMs, domains, etc.) using PTM-POSE and the ProteomeScoutAPI [45].

### Cell culture

The HepG2 cell line was obtained as a gift from the lab of Dr. Todd Bauer and LX293T cell line was obtained as a gift from the lab of Dr. Matthew Lazzara. Both cell lines were authenticated by Short Tandem Repeat (STR) profiling upon receiving. Cells were cultured in Dulbeco’s Modified Media (Gibco #11965092) and 10% FBS (Sigma Aldrich #F0926).

### Generating lentiviral vectors and particles

In order to best match ENCODE RBP knockdown experiments, we obtained bacterial glycerol stocks of validated MISSION shRNA plasmids matching shRNA hairpin sequences targeting two RBPs (U2AF1, SRSF3) from The RNAi Consortium (TRC) shRNA library (MilliPore-Sigma #SHCLNG, U2AF1 clone ID = TRCN0000001155, SRSF3 clone ID = TRCN0000273174). We confirmed the plasmid sequence for U2AF1-targeting plasmid by whole plasmid sequencing, but found an incorrect target sequence for the SRSF3-targeting plasmid we received. As a result, we cloned new hairpin sequences with the correct SRSF3 target sequence, as well as an additional plasmid containing hairpin sequences targeting GFP to serve as a control (GFP clone ID = TRCN0000072178). In brief, the original hairpin sequence was removed from the pLKO backbone by cutting with EcoR1 and Kpn1 restriction enzymes (New England Biolabs #R3101S, #R0142S). The pLKO backbone was isolated via gel electrophoresis and extracted with the Qiagen Gel Extraction Kit (Qiagen #28704). Two custom DNA oligonucleotides containing either the forward or reverse hairpin sequence were ordered from Thermo Fisher. Oligonucleotides were annealed using a thermocycler to heat them to 95°C and slowly decrease the temperature to room temperature over 70 cycles. The restricted backbone and annealed oligos were ligated using T4 DNA ligase (New England Biolabs #M0202) and the resulting plasmid was transformed into DH5*α* competent cells (Invitrogen #18265017). All hairpin sequences were validated by Sanger sequencing. Hairpin and target sequences are located in TableS6.

To produce shRNA-expressing lentiviral particles, LX293T cells and a second generation lentiviral packaging system was used. LX293T cells were transfected with packaging, envelope, and shRNA-expressing plasmids (3*µ*g pCMV-dR8.2, 0.5*µ*g pCMV-VSV-G, and 3*µ*g target, respectively) in 10 cm plates using FuGene6 (Promega #E269A). Virus was harvested and filtered through a 0.45 *µ*m filter (Cytiva #6780-2504) 36 and 60 hours post-transfection and stored at -70°C for future use.

### RBP knockdown, stimulation, and preparation of lysates

To knockdown GFP (control), U2AF1, or SRSF3, HepG2 cells were seeded in a 10 cm plate with 10 *µ*g/mL polybrene (Thomas Scientific #TR-1003-G) and 600 *µ*L of lentiviral particles. Viral amount was selected based on protein knockdown observed in a prior experiment. Media was replaced 17 hours after infection. 48 hours after infection, infected cells were passaged, seeded across 3 10 cm plates, and placed under 3 *µ*g/mL puromycin selection for five days. For the last 8 hours of selection, growth media was replaced with serum-free media in all plates to reduce basal phosphorylation levels. For one plate, cells were stimulated at the end of selection with 20 ng/mL EGF (Peprotech #AF-100-15), 20 ng/mL HGF (Genscript #Z03229), and 20 ng/mL insulin (Sigma-Aldrich #I6634) for five minutes prior to lysis. Serum starved and stimulated cells were lysed in 8M Urea (Sigma-Aldrich #U5378) supplemented with protease inhibitors, phosphatase inhibitors, and PMSF for downstream western blot and phosphoproteomic analysis. Protein concentration was quantified using a Pierce BCA protein assay (Thermo Fisher Scientific #23209).

### Western blot analysis

To assess protein-level impacts of RBP knockdown, protein and phosphorylation abundance was measured by western blot analysis. Protein lysates were reduced in Laemmli loading buffer (Boston BioProducts #BP-111R) and boiled for 10 minutes at 95°C. Standard SDS-PAGE analysis was performed using pre-cast NuPage 4-12% Bis-Tris gel, with one well containing a pre-stained protein ladder (Chameleon Duo) and the remaining wells loaded with equal amounts of total protein. Protein was transferred to a nitrocellulose membrane. Total protein transferred was visualized and quantified using Revert 700 total protein stain (LI-COR #926-11016) according to manufacturers instructions. Membranes were then blocked using Intercept Blocking Buffer (LI-COR #927-60001) diluted 1:1 in Tris Buffered Saline (TBS). After blocking, membranes were incubated with primary antibody solutions either overnight at 4°C or at room temperature for 1 hour, then washed with TBS-T (1% tween solution). After primary incubation, membranes were incubated with secondary antibody solutions for 1 hour at room temperature. Membranes were imaged with the LI-COR Odyssey DLx system. Membranes were stripped for reprobing using 0.2M NaOH. To quantify signals from each protein measurement, we used the ‘Quantitative Western Blot’ pipeline in Empiria Studio, normalizing band intensities to total protein signal from the Revert 700 total protein stain. Finally, for measurements of phosphorylated AKT or ERK, we normalized the phosphorylated signal relative to the total AKT or ERK signal, respectively. All primary and secondary antibodies and their dilutions utilized in this study can be found in TableS6.

### Sample preparation for mass spectrometry (MS)

After confirming knockdown of SRSF3 or U2AF1 in the corresponding lysates, we proceeded to phosphoproteomic analysis with mass spectrometry. In total, we analyzed a total of eighteen samples, with three biological replicates per sample group (shGFP, shU2AF1, and shSRSF3 either serum starved or stimulated). For the third MS replicate for SRSF3 knockdown lysates, we pooled two separate infection replicates into a single lysate stock in order to ensure sufficient protein for MS analysis. For each sample, 500 *µ*g of protein lysate was reduced with 5 mM TCEP (Thermo Fisher Scientific, #77720) for 15 min, alkylated with 10 mM iodoacetamide (MP Biomedicals, #100351) in the dark for 30 min, and then quenched with 10 mM DTT (Thermo Fisher Scientific, #R0861) for 15 min. Samples were digested with LysC (Fujifilm Biosciences Inc, #NC9242798) and Trypsin (Thermo Fisher Scientific, #90305) for 16 h at 37°C with shaking at 1000 rpm in a thermomixer. Peptides were desalted using Sep-Pak C18 cartridges (Waters, C18 Classic Cartridge, #WAT054925).

Phosphopeptide enrichment was performed on a KingFisher Flex system (Thermo Fisher Scientific) using MagReSyn zirconium-based immobilized metal affinity chromatography beads (Zr-IMAC HP; ReSyn Biosciences, #MR-ZHP005) according to a published protocol [46]. Briefly, peptides were resuspended in 200 *µ*L loading buffer (80% acetonitrile, 5% trifluoroacetic acid, and 0.1 M glycolic acid) and processed on the KingFisher using plates containing loading, wash, and elution buffers. For each sample, 33 *µ*L of Zr-IMAC HP beads suspended in 500 *µ*L acetonitrile were used. Beads were equilibrated in loading buffer, incubated with samples to bind phosphopeptides, and washed sequentially with loading buffer, wash buffer 2 (80% acetonitrile, 1% trifluoroacetic acid), and wash buffer 3 (10% acetonitrile, 0.2% trifluoroacetic acid). Bound phosphopeptides were eluted with 200 *µ*L of 1% NH4OH. Eluates were acidified and desalted using StageTips prior to drying and LC–MS/MS analysis.

### LC-MS/MS analysis

Peptides were separated on a Vanquish Neo ultra high-performance liquid chromatography (UHPLC) system configured in trap-and-elute mode using an Aurora Frontier TS C18 column (IonOpticks; 60 cm X 75 *µ*m inner diameter, 1.7 *µ*m particle size). Mobile phase A was water with 0.1% formic acid, and mobile phase B was acetonitrile with 0.1% formic acid. Peptides were eluted with a gradient from 1.0% to 9.6% B over 0.0–0.1 min at 0.3 *µ*L/min, from 9.6% to 14.4% B over 0.1–8.0 min at 0.3 *µ*L/min, and from 14.4% to 36.0% B over 8.0–50.0 min at 0.2 *µ*L/min. This was followed by a column wash at 85% B from 50.1 to 85.0 min at 0.3 *µ*L/min and re-equilibration for 4 column volumes at 2 *mu*L/min.

Data were acquired on an Orbitrap Astral mass spectrometer in DIA mode. Full MS scans were acquired at 240,000 resolution over an m/z range of 380–980 with AGC set to 250%. DIA MS2 scans were acquired at 80,000 resolution using 2 m/z isolation windows over an m/z range of 150–2000, with a maximum injection time of 3.5 ms, HCD collision energy of 25%, and 500% AGC.

### MS data analysis

DIA phosphoproteomics raw files were analyzed in Spectronaut v20.1.250624.92449 using the BGS Phospho PTM Workflow with the directDIA+ (Deep) workflow. Searches were performed against the UniProt humandatabase (downloaded in December 2024, 83,401 protein entries). PSM, peptide, and protein-group identifications were filtered at 1% FDR, with a precursor q-value cutoff of 0.01 and a phosphosite localization probability cutoff of 0.75. Quantification was based on MS2 peak area with cross-run normalization and interference correction enabled.

To compare quantification across replicates and ensure replicate consistency, we peformed principal components analysis (PCA). For this analysis, we restricted the data to peptides measured in all replicates and then normalized quantification across replicates through z-score standardization. Principal components analysis was then performed with the scikit-learn python package [47].

For each sample group, we calculated the average quantification across replicates for peptides measured in at least two of the three biological replicates and with a coefficient of variation (CV) less than 1. To identify peptides with similar quantification across sample groups, we performed K-means clustering with the scikit-learn python package. Using sample group means, we performed K-means clustering with k=3, then repeated clustering for each found cluster with K=2 to find subclusters. For comparison between sample groups (stimulation versus serum starvation, for example), log fold changes were calculated and statistical significance was assessed with a two-tailed t-test and Benjamini-Hochberg FDR correction.

### Gene set and phosphosite annotation enrichment

Gene set enrichment analysis was performed in python using the EnrichR module of the gseapy python package [48, 49]. All genes with differential phosphorylation in our group of interest (SRSF3-regulated genes, for example) were considered the foreground and all genes measured in the MS dataset were considered the background. We used Reactome gene sets for encrichment [23].

To identify functions associated with specific phosphorylation sites, we used function and biological process annotations from PhosphoSitePlus [22]. We used a hypergeometric test to calculate statistical enrichment, using the sites present in our group of interest (SRSF3-related sites, for example) as the foreground and all sites measured in the MS dataset as the background.

### Kinase Activity Analysis

Kinase activity analysis was performed with three different approaches – KSEA [20], KinaseLibrary [21], and KSTAR [19]. KSEA z-scores and false discovery rates were calculated based known kinase-substrates from PhosphoSitePlus [22] using a custom python-implementation of the algorithm described in the original publication [20]. KinaseLibrary enrichment scores were calculated using the ‘DiffPhosData’ class in the ‘kinase-library’ python package, using a p-value threshold of 0.05 and log fold change threshold of 2. Kinases with enrichment among downregulated and upregulated sites were not considerd. Finally, KSTAR (v1.1.0) activity predictions were generated using either up and down-regulated phosphosites as evidence of activity (*Log*2(*FoldChange*) ≥ 1,*FDR* ≤ 0.05) using the default parameters.

To create an aggregated ranking of the most up- and downregulated kinase in each group, we first ranked kinases in each algorithm individually. For KSEA, we used the Z-scores. For KinaseLibrary, we used the difference between the up- and down-regulated scores (*Change* = − *log*10(*p* − *adj*_*up*_) − *log*10(*p* − *adj*_*down*_)). Similarly, for KSTAR, we used the difference between activity scores among the up- and down-regulated sites. Once algorithm-specific ranks were calculated, we scaled these ranks between 0 and 1 (to account for the different number of kinases each algorithm generates predictions for) and averaged the ranks across the three algorithms. We then generated a final sorting of the kinases based on these ranks as can be seen in Figure S9.

### Predicting Splice Factor Activity with KATMAP

To identify splice factors relevant to the splice events observed after SRSF3 and U2AF1 knockdown, we harnessed a recently developed tool called KATMAP which relies on splicing perturbation information (such as ENCODE knockdown experiments) and RBP position-specific weight matrices (PWMs) [34]. For this analysis, we focused on RBPs with 1) knockdown RNA-sequencing data from ENCODE and that were either annotated as being splicing-related in the original ENCODE paper [13] or were annotated as being related to ‘mRNA splicing’ in UniProt [43]. Splice events were quantified from ENCODE knockdown experiments with rMATS-turbo as described in a prior section. We first obtained PWMs either from the KATMAP publication or from CISBP-RNA [35], prioritizing PWMs generated with direct measurements from RNAcompete or RNA-bind-N-seq experiments. For each RBP with an available PWM and quantified splice events, we generated a splicing model using KATMAP with default parameters, focusing on skipped exon events.

Before using the generated KATMAP models, we checked for good model fit as described in the original publication (*Zscore* ≥ 2, at least 2 coefficients are significantly non-zero). In total, this resulted in models for 27 different RBPs. We applied each of these models to splice events observed after SRSF3 and U2AF1 knockdown to identify RBPs with the largest influence on the observed splicing changes. Finally compared the applied model results to changes in transcript splicing or expression (from ENCODE) or changes in phosphorylation from MS.

### Statistical analysis

Unless otherwise stated, all data and statistical analysis performed in this work was done in python using the SciPy package [50], Scikit-learn package [47], or custom-built functions. P-values were corrected using Benjamini-Hochberg false discovery rate correction.

## Code and Data Availability

The raw mass spectrometry and metadata associated with this study has been deposited to the ProteomeXchange Consortium (http://proteomecentral.proteomexchange.org/) via the PRIDE partner repository [51] with dataset identifier PXD060364. Additional data associated with downstream analysis for this study, including downloading and processing of ENCODE knockdown experiments, have been deposited in Figshare (https://doi.org/10.6084/m9.figshare.32028282). All code used to process data and generate figures has been uploaded to a github repository (https://github.com/NaegleLab/SF_KD_Phosphoproteomics_Analysis).

## Acknowledgements

Research reported in this publication was supported by the National Institute Of General Medical Sciences of the National Institutes of Health under Award Number R35GM138127, by the National Cancer Institute (NCI) under Award Number U01CA284193, and by the National Science Foundation Graduate Research Fellowship under Grant No. 1842490. In addition, it was supported by a pilot award from the University of Virginia Comprehensive Cancer Center. T.Z. was also supported by NCI grant R00CA273170. The content is solely the responsibility of the authors and does not necessarily represent the official views of the National Institutes of Health or the National Science Foundation. The methods figure in Figure 1 contains icons obtained from Bioicons.com created by Simon Durr and Marcel Tisch. We would also like to think Yogi Raghav and John Platig for their helpful guidance when brainstorming this project and navigating ENCODE datasets.

## Declaration of generative AI and AI-assisted technologies

During preparation of this work, we used Claude.ai to identify grammatical errors in our writing and increase clarity of the manuscript. After using this tool, we reviewed the content and edited our original text as needed and take full responsibility for the content of the publication.

