## Supplementary Figures for "Evaluating splicing factor and kinase network crosstalk through global phosphoproteomics"

### Investigating the influence of splicing factors, SRSF3 and U2AF1, on the phosphoproteome

**1 University of Virginia, Department of Biomedical Engineering, Charlottesville, VA, 22903**

*
† authors contributed equally to this work

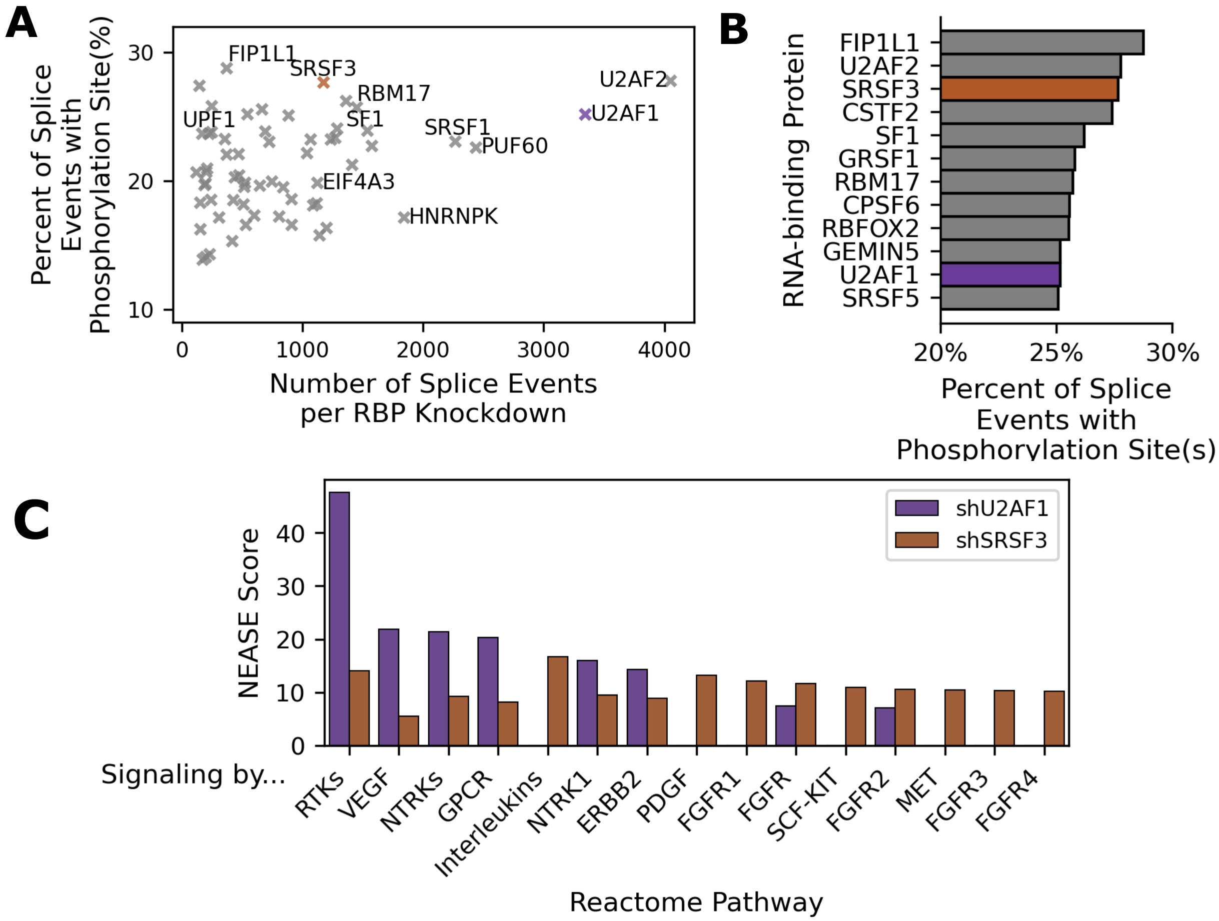

**Figure S1. Identifying Candidate Splice Factors for Phosphoproteomic Analysis** To select splice factors for phosphoproteomic analysis, we first reanalyzed RNA-binding protein (RBP) knockdown experiments from ENCODE to identify splicing-associated RBPs that drive many signaling-related splice events [1]. Due to a high number of splice events associated with phosphorylation sites and RTK signaling, we ultimately selected SRSF3 and U2AF1 for further analysis. **A)** Number of significant splice events ($\Delta PSI \geq 20\%$, $FDR \leq 0.05$) induced by RBP knockdown for all knockdown experiments analyzed (61 total). We compared this to the fraction of these events that impact phosphorylation sites. U2AF1 (purple) and SRSF3 (brown) are highlighted in color. **B)** Ranked list of RBPs based on the percent of knockdown-associated splice events containing phosphorylation sites, with U2AF1 and SRSF3 highlighted in color. **C)** NEASE enrichment scores for splice events associated with either U2AF1 or SRSF3 knockdown, focusing on Reactome gene sets denoted with "Signaling by ___" [2,3]

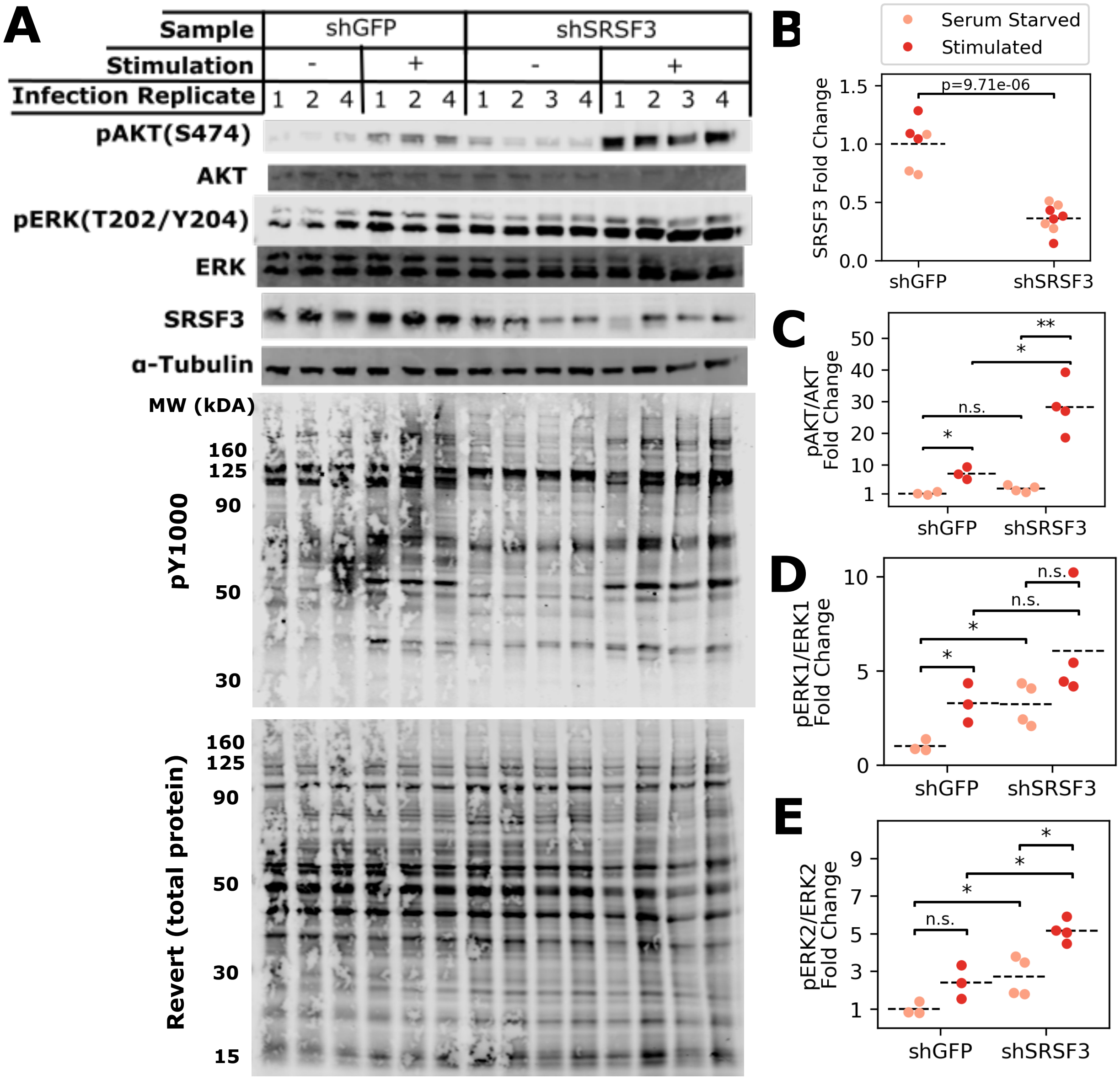

**Figure S2. Validation and western blot assessment of SRSF3 knockdown.** HepG2 cells were transduced with lentiviral plasmids expressing shRNA targeting either GFP or SRSF3 and selected with puromycin for 5 days. Selected cells were serum starved for 8 hours and stimulated with 20ng/mL of EGF, HGF, and insulin for 5 minutes. The resulting protein lysates were assessed by western blot, with 10$\mu$g protein loaded per well. Four infection replicates are shown for shSRSF3 and three infection replicates are shown for shGFP samples. **A)** Cropped western blot images showing probes targeting SRSF3, total and phosphorylated AKT (S473), total and phosphorylated ERK1/2 (T202/Y204), total pY signal (pY1000), and total protein (Revert stain). $\alpha$-Tubulin is shown as loading reference. **B-E)** Fold changes in band intensity quantified from western blot in Panel A, relative to average band intensity from shGFP control samples. Signal intensity was first normalized to either total protein (SRSF3) or individual protein signal (pERK1/2, pAKT). Signal intensity was quantified and normalized using Empiria Studio. Statistical significance was assessed using a two-tailed t-test and is shown on each plot (**:$p \leq 0.001$, *:$p \leq 0.05$, n.s.: not significant).

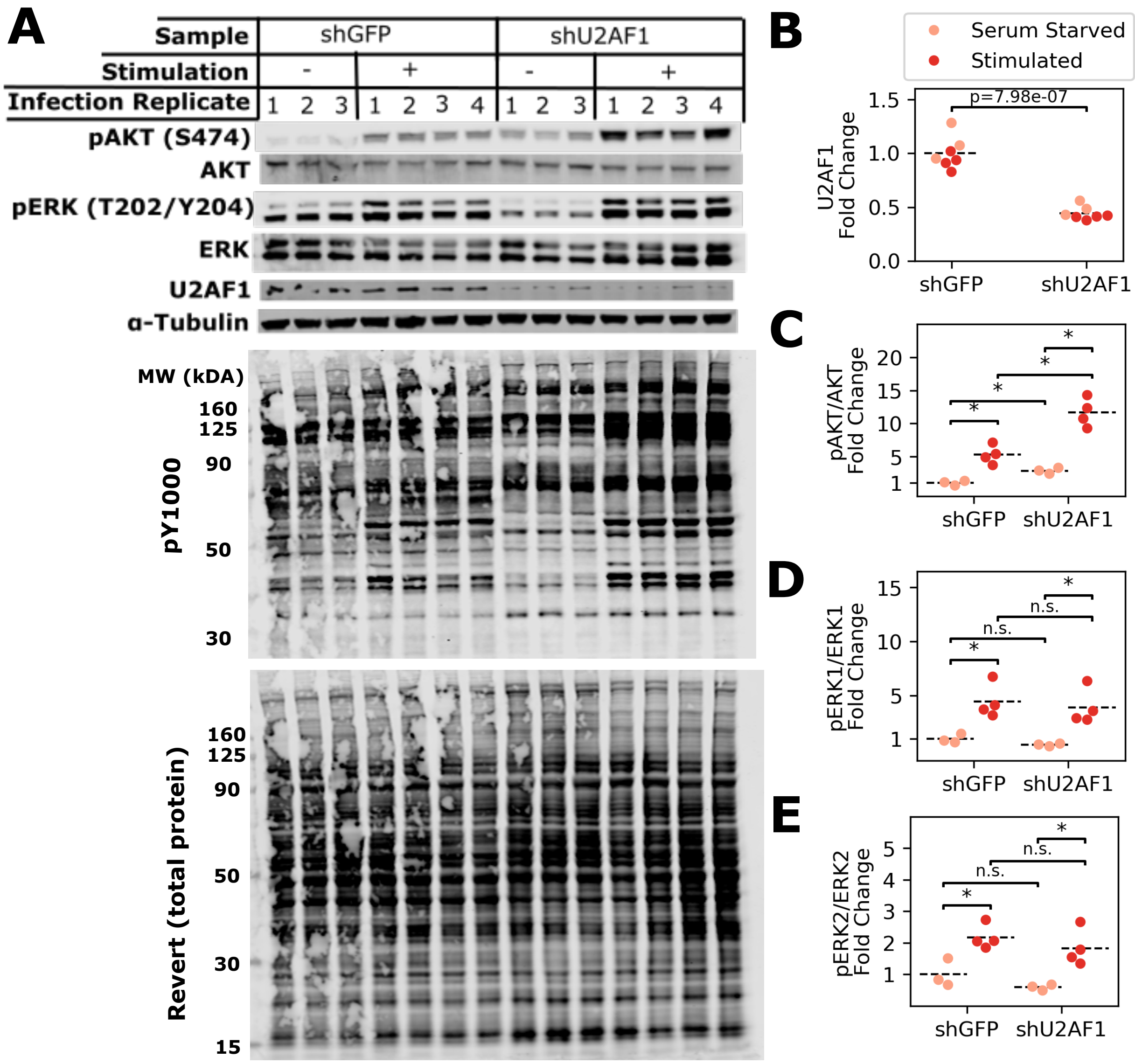

**Figure S3. Validation and western blot assessment of U2AF1 knockdown.** HepG2 cells were transduced with lentiviral plasmids expressing shRNA targeting either GFP or U2AF1 and selected with puromycin for 5 days. Selected cells were serum starved for 8 hours and stimulated with 20ng/mL of EGF, HGF, and insulin for 5 minutes. The resulting protein lysates were assessed by western blot, with 30$\mu$g protein loaded per well. For each shRNA target, four infection replicates are shown for stimulated cells and three infection replicates are shown for the serum starvation conditions. **A)** Cropped western blot images showing probes targeting U2AF1, total and phosphorylated AKT (S473), total and phosphorylated ERK1/2 (T202/Y204), total pY signal (pY1000), and total protein (Revert stain). $\alpha$-Tubulin is shown as loading reference. **B-E)** Fold changes in band intensity quantified from western blot in Panel A, relative to average band intensity in shGFP control samples. Signal intensity was first normalized to either total protein (U2AF1) or individual protein signal (pERK1/2, pAKT). Signal intensity was quantified and normalized using Empiria Studio. Statistical significance was assessed using a two-tailed t-test and is shown on each plot (**:$p \leq 0.001$, *:$p \leq 0.05$, n.s.: not significant).

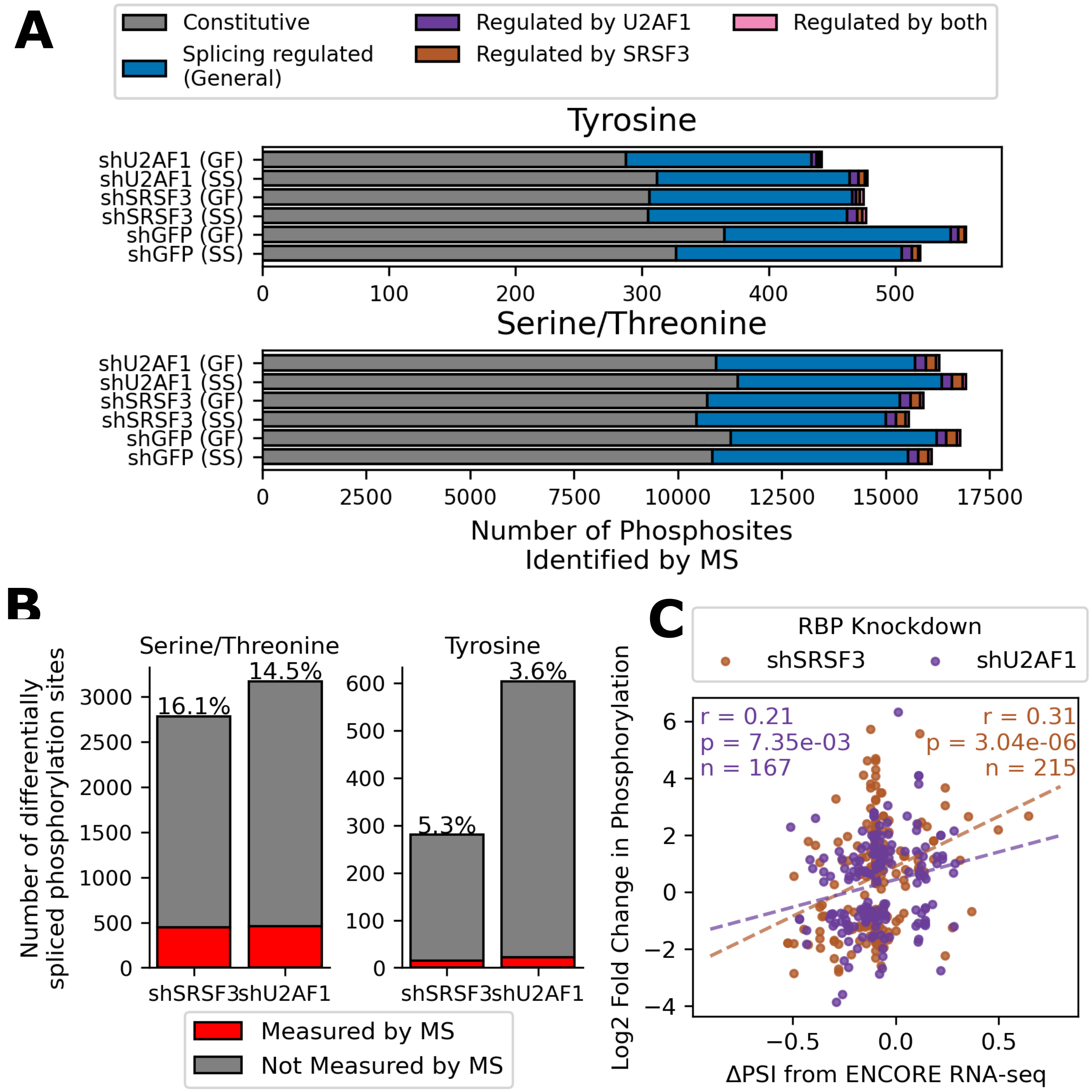

**Figure S4. Determining connection between differential inclusion of phosphorylation sites and measured phosphorylation** To determine if splicing of phosphorylation-encoding regions could help explain differential phosphorylation, we identified phosphorylation sites associated with splice events after knockdown (based on ENCODE data [1]) and compared them to phosphorylation site measurements from DIA-MS in this study. **A)** Number of phosphorylation sites identified in each sample (measured in at least 2 replicates). We broke down phosphorylation sites based on whether they were regulated by SRSF3 and/or U2AF1 knockdown based on ENCODE data, have previously been shown to be splicing regulated in other contexts, or are constitutive PTMs (always found in protein isoforms). Constitutive vs. non-constitutive sites identified using information from PTM-POSE [4]. **B)** Number of serine/threonine or tyrosine sites associated with alternative splicing events from ENCODE data, and the fraction of these sites that were also measured by mass spectrometry in this study. **C)** Comparison of the change in phosphorylation after SRSF3 (brown) or U2AF1 (purple) knockdown to change in percent spliced in (PSI) for significant splice events based on ENCODE data. Pearson correlation for each knockdown shown on the plot.

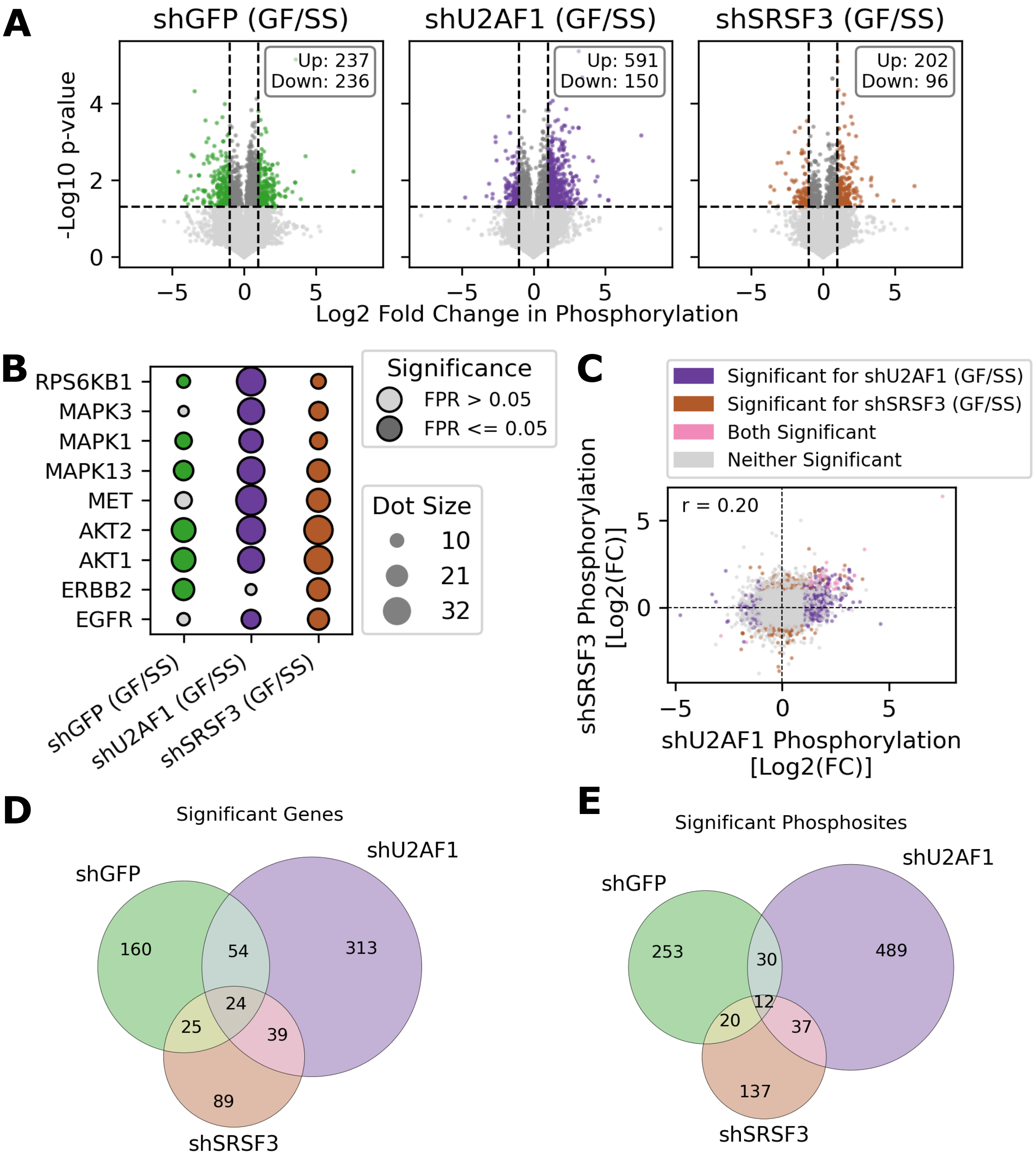

**Figure S5. Differential phosphorylation induced by stimulation across RBP knockdown conditions.** HepG2 cells expressing shGFP (control), shU2AF1, or shSRSF3 were serum starved (SS) and stimulated with a cocktail of growth factors (GF = EGF, HGF, and insulin) for 5 minutes to drive RTK activation. **A)** Fold change in phosphorylation after stimulation (GF/SS) and corresponding p-values (uncorrected) for each knockdown condition ($p \leq 0.05$, $|Log2(FC)| \geq 1$). **B)** Kinases with increased activity after stimulation, based on KSTAR predictions and focusing on kinases we would expect to be active after RTK activation [5]. Dotsize indicates degree of activity, and color indicates significance. **C)** Comparison of fold changes in phosphorylation for sites measured after SRSF3 and U2AF1 knockdowns. Color indicates whether phosphorylation changes were significant for shU2AF1 (purple), shSRSF3 (brown), or both (pink) ($p \leq 0.05$, $|Log2(FC)| \geq 1$). **D)** Overlap in the genes with differential phosphorylation events after stimulation in each knockdown condition. **E)** Overlap of the individual sites with differential phosphorylation events after stimulation in each knockdown

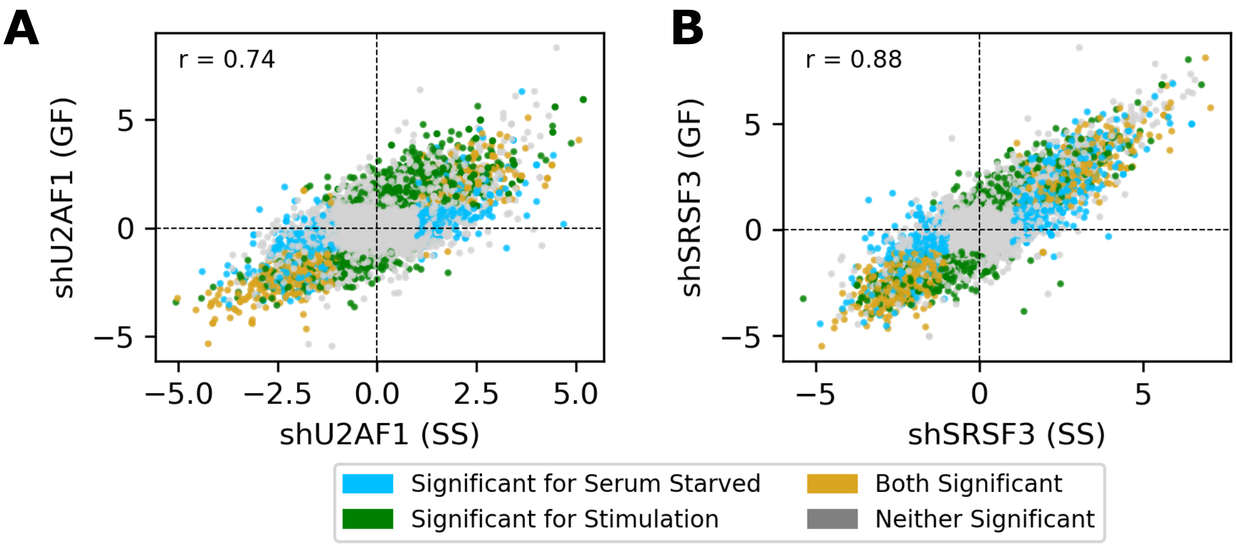

**Figure S6. Perturbations caused by knockdown are comparable across stimulation conditions.** To determine the importance of stimulation condition in defining the **(B)** U2AF1-associated phosphoproteome or the **(B)** SRSF3-associated phosphoproteome, we compared the fold changes in phosphorylation after knockdown relative to the shGFP control under either serum starvation or stimulation. In both panels, points are colored based on whether the fold change was found to be significant under serum starvation (blue), stimulation (green), or both (gold) ($p \leq 0.05$, $|Log2(FC)| \geq 1$). Pearson correlation is shown in the upper left hand of the plot.

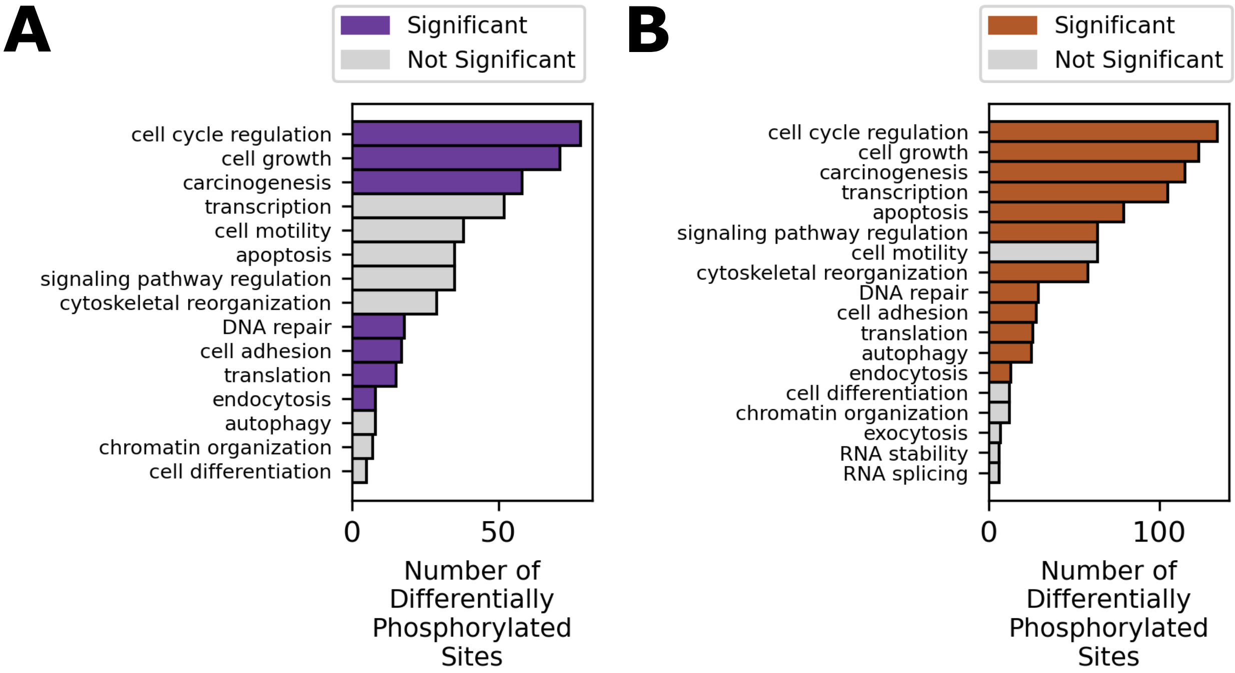

**Figure S7. Functions associated with phosphosites perturbed by U2AF1 and SRSF3 knockdown. A)** Biological processes associated with differentially phosphorylated sites after U2AF1 knockdown ($log2FC \leq -1$, $p_{adj} \leq 0.05$). Annotations based on PhosphoSitePlus [6]. Statistical enrichment was assessed with a hypergeometric test, with significant processes denoted in purple ($p \leq 0.05$) **B)** Biological processes associated with differentially phosphorylated sites after SRSF3 knockdown ($log2FC \leq -1$, $p_{adj} \leq 0.05$). Annotations based on PhosphoSitePlus [6]. Statistical enrichment was assessed with a hypergeometric test, with significant processes denoted in brown ($p \leq 0.05$)

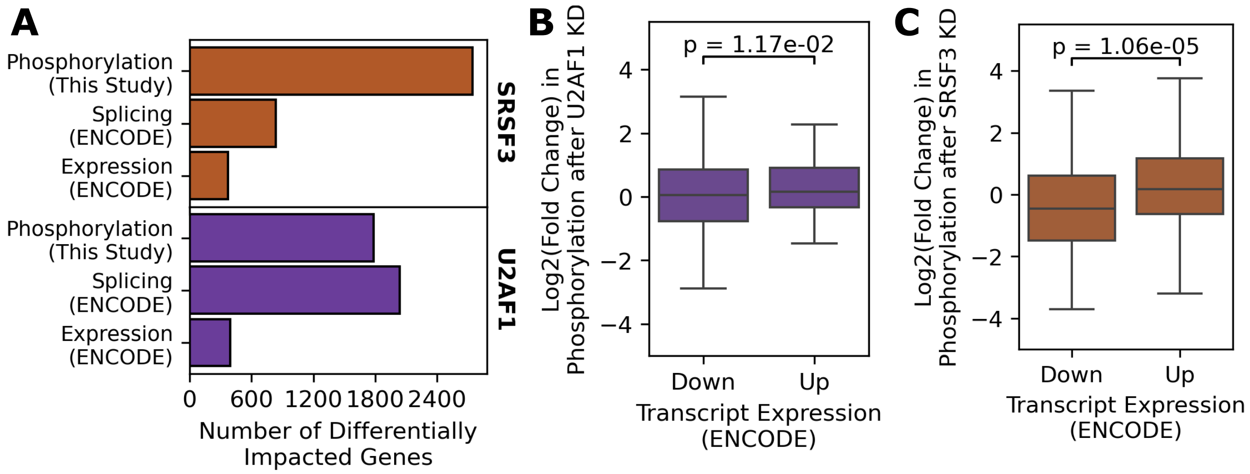

**Figure S8. Comparison of differentially phosphorylated genes and regulated transcripts.** To determine how mRNA processing facilitated by RBP knockdown may relate to differential phosphorylation events, we compared the genes identified as differentially expressed and/or spliced based on ENCODE data to genes observed to be differentially phosphorylated after SRSF3/U2AF1 knockdown [1]. **A)** Number of differentially phosphorylated, differentially expressed, or alternatively spliced genes after SRSF3 or U2AF1 knockdown. Expression/splicing information was based on RNA-sequencing data from ENCODE [1]. Significance was defined as $|log2FC| \geq 1$ and $p_{adj} \leq 0.05$ for differential expression and phosphorylation. Significance was defined as $\Delta PSI \geq 0.2$, $FDR \leq 0.05$, and a minimum of 20 read counts for alternative splicing events. **B)** Fold changes in phosphorylation for differentially expressed genes after U2AF1 knockdown. Upregulated transcripts tended to have higher phosphorylation fold changes based on a t-test ($p \leq 0.05$).**C)** Fold changes in phosphorylation for differentially expressed genes after SRSF3 knockdown. Upregulated transcripts tended to have higher phosphorylation fold changes based on a t-test ($p \leq 0.05$).

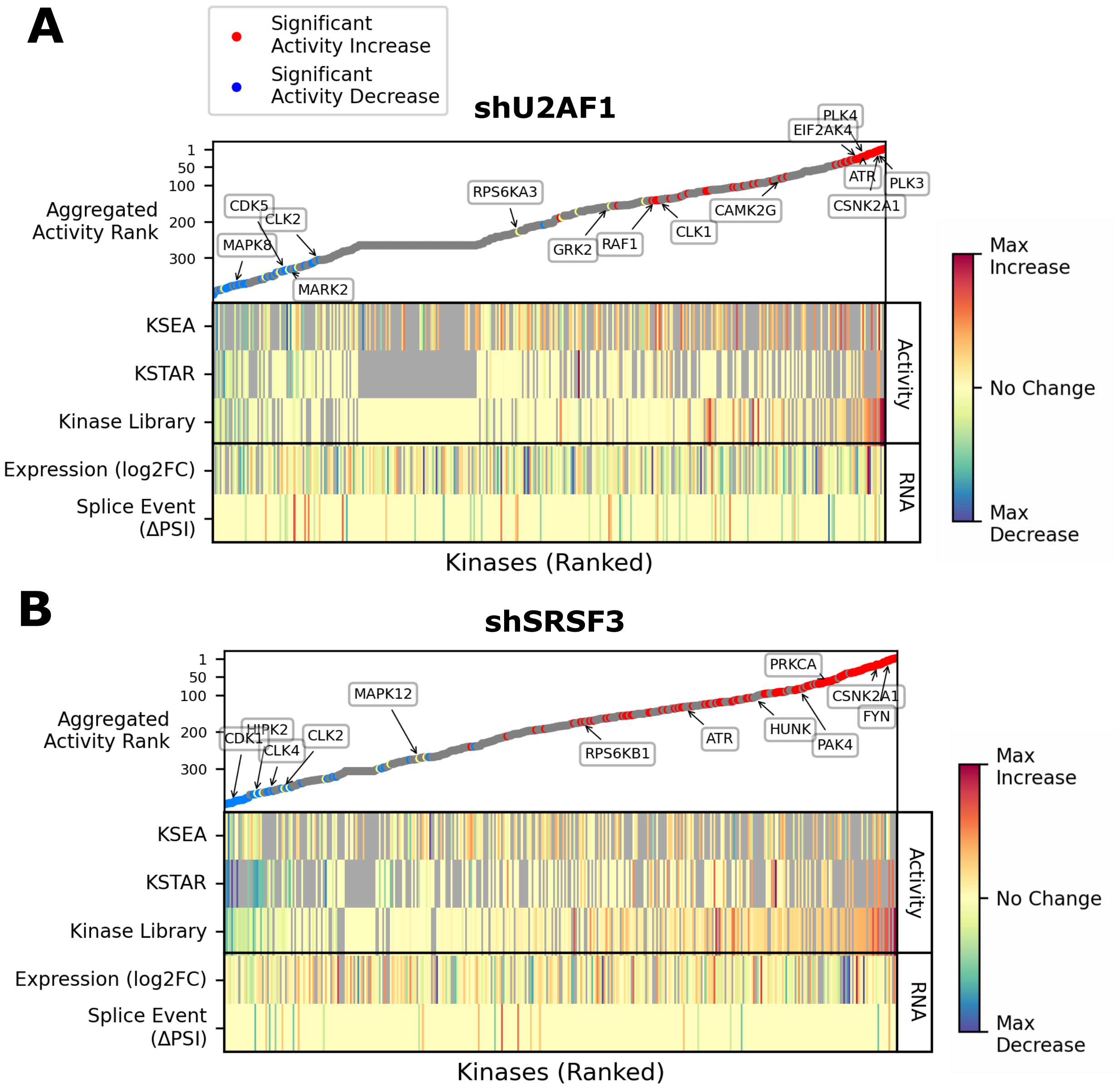

**Figure S9. Comparing kinase activity changes to mRNA regulation by SRSF3 and U2AF1.** Ranked kinase activity changes and associated transcript regulation (expression and splicing changes based on ENCODE data [1]). We did this analysis for **A)** U2AF1 knockdown and **B)** SRSF3 knockdown. Kinase activity was predicted using three approaches – KSEA [7], KinaseLibrary [8], and KSTAR [4]. For each algorithm, the change in activity after knockdown was calculated for each kinase and kinases were ranked from largest increase to largest decrease in activity. Ranks were aggregated across all three algorithms to obtain a final activity rank, shown in the top scatter plot (see methods for more details). Kinases were then colored based on whether the kinase exhibited a significant activity change in at least one algorithm (blue = significant decrease, red = significant increase). Shown below this plot are results from individual activity algorithms, scaled to be on the use the same color bar. We also compared to any changes in exon inclusion or mRNA expression based on ENCODE information [1]. Exon inclusion was assessed by rMATS [9], with only the most significant event being shown for each kinase (min = -0.8, max = 0.8). mRNA expression was based on Log2 fold changes (min = -3, max = 3). Some kinases of interest, either due to particular pathway or because it is regulated at the RNA level, are labeled on the plot. Full results can be found in Supplementary Table 4.

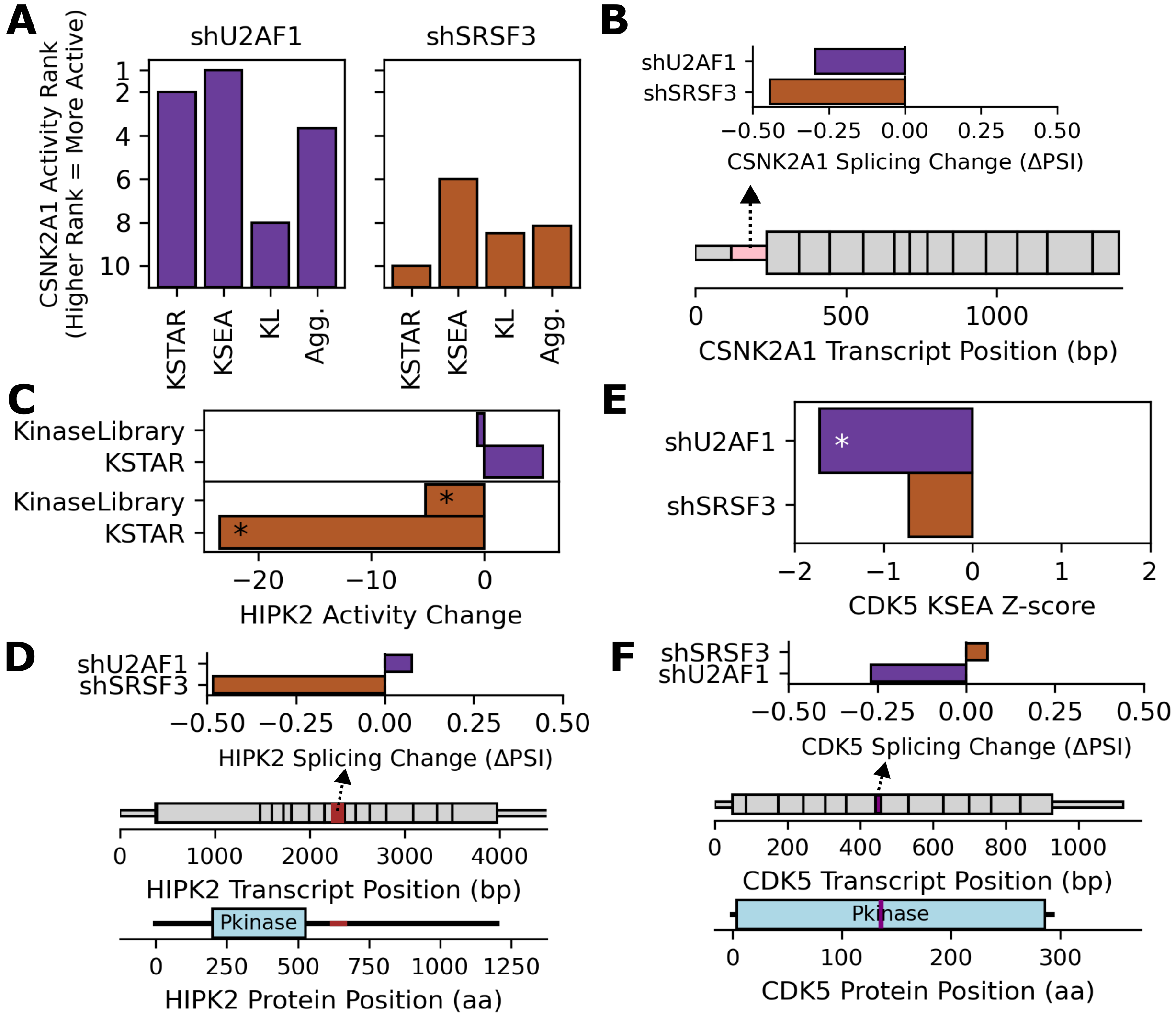

**Figure S10. Examples of kinases with activity influenced by alternative splicing.** We identified several kinases with differential activity after SRSF3 (brown) or U2AF1 (purple) knockdown, which were also alternatively spliced based on ENCODE and rMATS activity [1,9]. Activity predictions were based on KSEA [7], KinaseLibrary (KL) [8], and/or KSTAR [4]. **A)** Kinase activity rank for CSNK2A1 after SRSF3 or U2AF1 knockdown. Higher rank (smaller value) indicates large increase in activity, lower rank (larger value) indicates large decrease in activity. Ranks based on individual activity algorithms or aggregated across each algorithm. **B)** Skipped exon event occurring in the noncoding region of CSNK2A1 and the change in percent spliced in (ΔPSI) after SRSF3 and U2AF1 knockdown. The exons and protein domains associated with CSNK2A1 are shown with the splice event region highlighted in pink. **C)** HIPK2 activity changes based on differences in KSTAR or KinaseLibrary scores among up- and down-regulated phosphorylation sites. *: $p_{adj}$ *or* $FPR \leq 0.05$. **D)** 3' alternative splice site event in HIPK2 occurring after SRSF3 knockdown and the change in percent spliced in (ΔPSI) after SRSF3 and U2AF1 knockdown. The exons and protein domains associated with HIPK2 are shown with the splice event region highlighted in brown. The splice event was previously validated after SRSF3 knockdown and affects a region previously identified as an E3 ligase binding region [10]. **E)** CDK5 activity change after knockdown, based on KSEA z-scores. *: $p \leq 0.05$. **F)** Skipped exon event occurring within CDK5 kinase domain and the change in percent spliced in (ΔPSI) after SRSF3 and U2AF1 knockdown. The exons and protein domains associated with CDK5 are shown with the splice event region highlighted in purple.

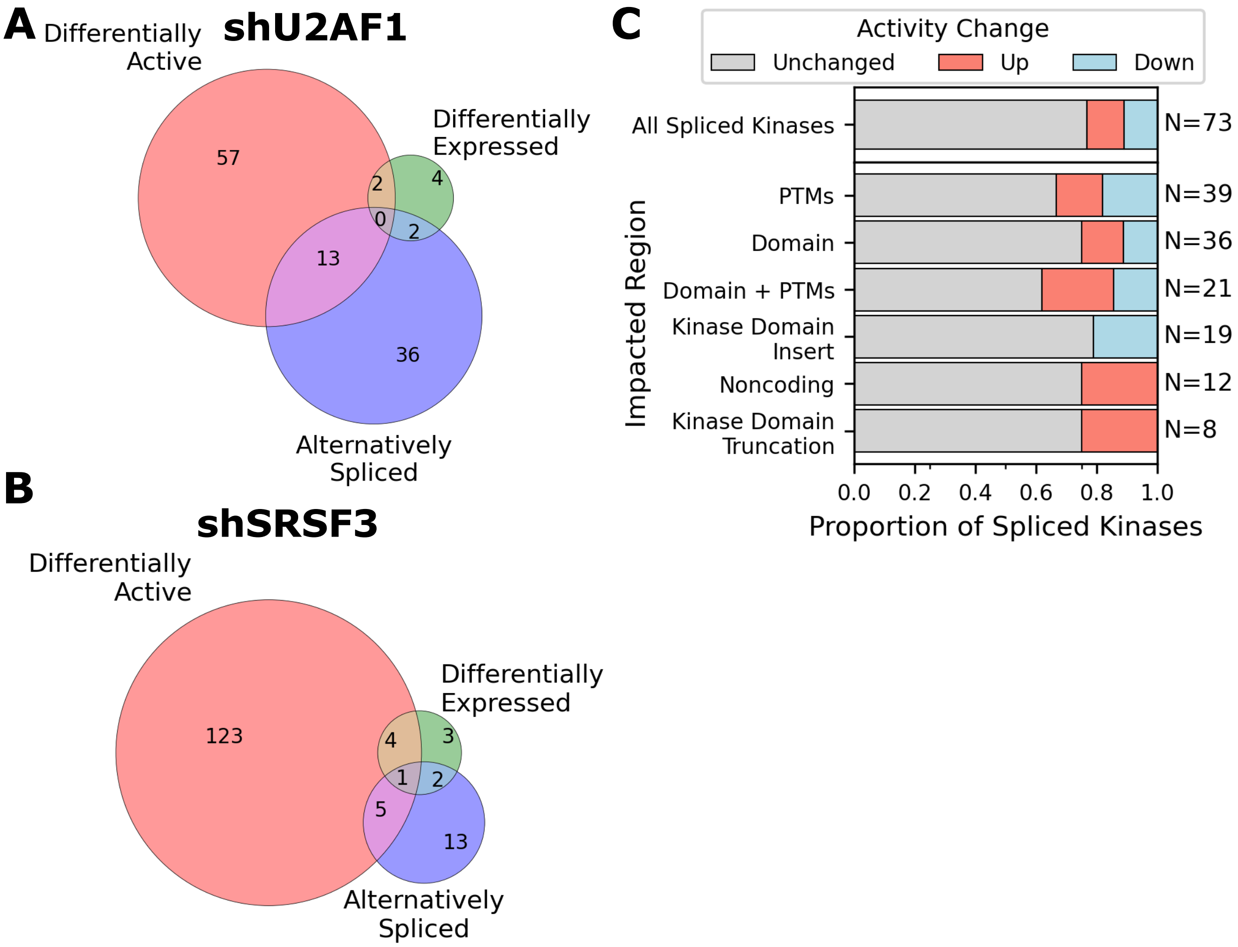

**Figure S11. Overlap between kinase activity changes between knockdowns.** Kinase activity changes were predicted using KSEA [7], KinaseLibrary [8], and KSTAR [5]. We then compared these activity perturbations across knockdowns and to transcript regulation of the same kinases **A)** Number of kinases with differential kinase activity after U2AF1 knockdown that are also regulated through differential mRNA expression or alternative splicing, based on ENCODE data. **B)** Number of kinases with differential kinase activity after U2AF1 knockdown that are also regulated through differential mRNA expression or alternative splicing, based on ENCODE data. **C)** Proportion of kinases undergoing a splice event that had a change in activity (up = red, down = blue). We repeated this analysis for splice events impacting specific features of the transcript/protein, including the noncoding region, protein domains, and PTMs. "Kinase domain insert" refers to differential inclusion of an exon that resides in the middle of the kinase domain. "Kinase Domain Truncation" refers to exons that exist on the edge of the kinase domain (partially overlap it). The number of kinases associated with each category of splice event are indicated on the right of each bar.

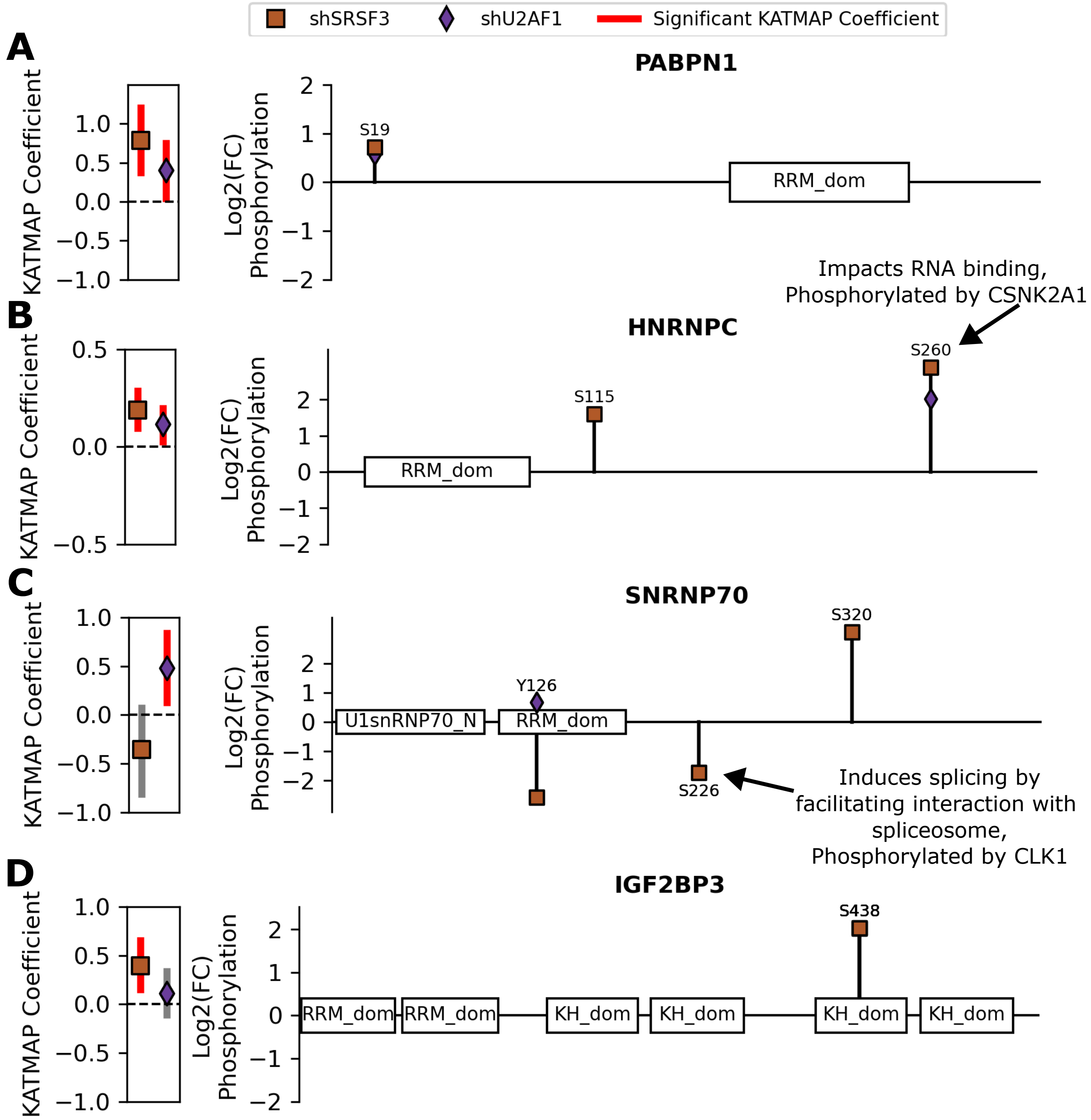

**Figure S12. Examples of phosphorylation regulation of splice factors and their activity.** After identifying altered splice factor activity after SRSF3 and U2AF1 knockdown based on KATMAP models and ENCODE data [1,11], we sought to determine if phosphorylation could be connected to the change in splice factor behavior. We noted many different examples of phosphorylation regulation of splice factors, including **A)** PABPN1, **B)** HNRNPC, **C)** SNRNP70/U1-70K, and **D)** IGF2PB3. For each, we show the KATMAP coefficients in the left panel and whether there was a significant change (94% confidence interval doesn't overlap with 0). In the right panel, we show the protein architecture (length and domains) with the Log2 fold change in phosphorylation for any sites in the splice factor that were significantly changed ($p_{adj} \leq 0.05$). We also annotated sites with known function and/or kinases on the plot itself. For both panels, data from SRSF3 knockdown is shown as a brown square, while data from U2AF1 knockdown are shown as a purple diamond.
